# TMEM55B promotes anterograde transport and distal axonal distribution of LAMP1-positive organelles

**DOI:** 10.64898/2026.10.27.762112

**Authors:** Sönke Rudnik, Chun Hei Li, Dieter-Christian Gottschling, Alexander Veh, Daniel J. Barbosa, Julia Richter, Stijn Stroobants, Björn Schuster, Inken M. Beck, Radislav Sedlack, Robert Häsler, Mara Riechmann, Frank Döring, Paul Saftig, Rudi D’Hooge, Katsuji Yoshioka, Reto Gassmann, Ginny G. Farias, Markus Damme

**Author notes:** Corresponding author: Prof. Dr. Markus Damme, Faculty of Chemistry, Biochemistry I (BCI), University of Bielefeld, Bielefeld, Germany. these authors contributed equally (Sönke Rudnik, Chun Hei Li, Dieter-Christian Gottschling, Alexander Veh).

## Abstract

Lysosomal function throughout life is vital for the proper functioning and survival of post-mitotic neurons by mediating cellular signaling, degradation, and recycling processes. Their intracellular transport relies on microtubule-dependent mechanisms involving kinesin and dynein motors. While lysosomes were long thought absent from axons, recent studies have demonstrated the importance of LAMP1-positive endo-/lysosomal organelles in axonal compartments. This study investigates the physiological role of TMEM55B, a lysosomal membrane protein involved in lysosome positioning and motor coupling in neurons. TMEM55B is highly expressed in LAMP1-positive organelles in cortical neurons. We generated *viable Tmem55b* knockout mice that exhibited altered lysosomal positioning in neurons. Lysosomes clustered abnormally in the pre-axonal exclusion zone and the proximal axon, with many forming elongated tubular structures. Similarly, in primary rat neurons, TMEM55B depletion resulted in lysosomal tubulation and impaired bidirectional lysosomal transport, particularly in distal axon segments, without affecting lysosome velocity in the proximal and mid-axon. These transport defects were consistent with disrupted JIP4 function, a scaffold linking TMEM55B to both dynein and kinesin motors. TMEM55B deficiency did not result in neurodegeneration, behavioral deficits, or reduced lifespan in mice. As an orthogonal approach, we analyzed TMEM55B function in *C. elegans*. Worms lacking the TMEM55B homolog Y71H2AM.2 showed altered lysosomal dynamics in cholinergic neurons, supporting an evolutionarily conserved role. To test the involvement of TMEM55B’s paralog, TMEM55A, in regulating neuronal lysosome positioning, we generated *Tmem55a/b* double-knockout mice, which exhibit impaired growth, premature death, and an overall aggravated phenotype. Collectively, our findings demonstrate that both TMEM55B and TMEM55A regulate lysosome distribution in neurons, particularly in axons, by influencing JIP4-dependent retro- and, notably, anterograde motor protein-mediated transport. This function is critical for spatial lysosomal homeostasis. Our study underscores the importance of TMEM55A/TMEM55B in regulating neuronal organelle dynamics and may be a factor considered in neurodegenerative diseases involving lysosomal trafficking defects.

## Introduction

Lysosomes are critical organelles for degrading intra- and extracellular macromolecules taken up by endocytosis or autophagy, respectively. They are essential for nutrient recycling and for removing protein aggregates and defective organelles in eukaryotic cells by mediating the final steps of autophagy. Their function is vital for the proper function and survival of post-mitotic neurons, and, not surprisingly, dysregulation of lysosomal degradative pathways is linked to various neurodegenerative disorders (Wallings et al., 2019; Wang et al., 2018). Therefore, a comprehensive understanding of lysosomal functions and their impact on neuronal viability and function is essential to clarify their role in health and disease. Lysosomes are highly dynamic organelles, and their intracellular transport is mediated via microtubule tracks over longer distances. Their spatio-temporal distribution depends on active transport driven by the anterograde- and retrograde motors kinesin and dynein, respectively (Bonifacino and Neefjes, 2017; Saftig and Puertollano, 2021). In neurons, the spatial distribution of lysosomes, late endosomes (LE), and related organelles containing the prototypical marker LAMP1 is critical due to the highly complex specialization of the somatodendritic and axonal compartments in which these organelles fulfill different functions. Although the axon was formerly thought not to contain lysosomes at all, and the great majority of the total pool of lysosomes in neurons is localized in the somatodendritic compartment (Becker et al., 1960), more recent work shows compelling evidence for an important physiological function of lysosomes or related LAMP1^+^-organelles in the axon (Adnan et al., 2020; Bonifacino and Neefjes, 2017; Cason et al., 2022; Cheng et al., 2018; De Pace et al., 2020; Farfel-Becker et al., 2019; Farias et al., 2015; Farias et al., 2017a; Roney et al., 2022; Winckler et al., 2018). The relevance of axonal endo-/lysosomal organelles is further highlighted by the fact that they pathologically accumulate in axonal spheroids. Axonal swellings are found, e.g., as a prominent feature of amyloid plaques in Alzheimer’s disease (Cataldo and Nixon, 1990; Lie et al., 2021).

Axonal LAMP1^+^-organelles might not fulfill all requirements for the definition of lysosomes, i.e., they often contain fewer or no lysosomal hydrolases (Cheng et al., 2018; Lie et al., 2021; Winckler et al., 2018; Yap et al., 2018), but for simplicity, we will refer to them as lysosomes. Lysosomes fuse with autophagosome in the distal axon, and they mature on the way back to the soma (Maday et al., 2012). The entry of cargo (including lysosomes) into the axon at the proximal axon and the axon initial segment (AIS) is tightly regulated, and the number and composition of lysosomes differ between the proximal axon and the distal axon (Farias et al., 2017b; Ozkan et al., 2021).

While an actin-based mesh was previously thought to regulate cargo entry into the AIS (Song et al., 2009), more recent work, however, suggests that the region proximal to the AIS, the so-called pre-axonal exclusion zone (PAEZ), governs cargo entry into the axon, including lysosomes (Farias et al., 2015). A model was proposed in which polarized cargo sorting at the PAEZ is microtubule-based and depends on the differential acquisition of microtubule motors that mediate selective transport into the axon (Farias et al., 2015). Consequently, only cargo bound to kinesins can pass the PAEZ (Farias et al., 2015).

Different motor protein complexes mediate the microtubule-dependent transport of lysosomal cargo vesicles: Kinesin-1 and kinesin-3 are the principal anterograde motors mediating the transport of lysosomes (Guardia et al., 2016). Both kinesins depend, among other alternative adaptor machineries, on the cytosolic BLOC-one-related complex (BORC) (Cason and Holzbaur, 2022; Guardia et al., 2016). BORC interacts with the Ragulator complex, which is attached to the lysosomal membrane by lipidation (Pu et al., 2017; Pu et al., 2015). ARL8B, a small GTPase, recruits the BORC complex and its effector SKIP/PLEKHM2, which directly recruits kinesin-1. Alternative adaptor mechanisms on lysosomes for kinesins have been described, among the scaffolds c-Jun NH2-terminal kinase (JNK)-interacting protein 3 (JIP3) and JIP4 (Vilela et al., 2026; Watt et al., 2015). The retrograde transport of lysosomes is mainly mediated by the complex of dynein/dynactin. A complex composed of the small GTPase RAB7 and Rab-interacting lysosomal protein (RILP) couples lysosomes to the dynein/dynactin complex (Ballabio and Bonifacino, 2020; Cason and Holzbaur, 2022).

More recently, a RAB7/RILP-independent mechanism mediating the retrograde transport of lysosomes was identified: TMEM55B, an integral lysosomal membrane protein with two transmembrane segments, a large N-terminal cytoplasmic domain, and a short cytoplasmic C-terminus, interacts with JIP4. JIP4 binds dynein/dynactin and mediates retrograde lysosomal transport (Willett et al., 2017). Overexpression of TMEM55B in ARPE19 or HeLa cells leads to perinuclear clustering of lysosomes, and KD drives microtubule-dependent transport of lysosomes to the cell’s periphery. Likewise, the KD of JIP4 drives lysosomes into the cell’s periphery (Willett et al., 2017). JIP4 is a large scaffold that binds both dynein and, notably, also kinesin-1, which mediates anterograde organelle transport (Celestino et al., 2022; Montagnac et al., 2009; Nguyen et al., 2005; Verhey et al., 2001; Vilela et al., 2019). Accordingly, JIP3 (a paralog of JIP4) can induce kinesin activity *in vitro* (Vilela et al., 2026; Watt et al., 2015), and JIP3/4 can also activate dynein (Cason and Holzbaur, 2023). Binding of either dynein or kinesin-1 to JIPs is not exclusive and is non-competitive (Vilela et al., 2026).

TMEM55B can undergo reversible phosphorylation and is S-palmitoylated (Rudnik et al., 2022; Takemasu et al., 2019), suggesting a complex regulation of the protein’s function by posttranslational modifications. Besides JIP4, TMEM55B binds additional interaction partners (mostly regulating endo-/ lysosomal function), suggesting that it may act more generally as a scaffold (Damme and Kummel, 2026; Waschbusch et al., 2026). Among these is TBC1D9B, a GTPase-activating protein (GAP) for which we have recently shown that it inactivates ARL8B, providing an additional layer of complexity for TMEM55B regulating lysosomal positioning (Duhay et al., 2026).

A paralog of TMEM55B in eukaryotes is TMEM55A, which shares a high degree of similarity and suggests similar function(s). Notably, both are localized in late endosomes and lysosomes (Ungewickell et al., 2005). TMEM55A and TMEM55B were initially identified as phosphatidylinositol (PI)(4,5)P_2_ 4-phosphatase (Ungewickell et al., 2005). This function was later challenged (Pal et al., 2023; Willett et al., 2017), and phosphatase activity could not be reproduced. More recent studies, however, support a PI(4,5)P_2_ 4-phosphatase function for TMEM55A (Liu et al., 2025; Morioka et al., 2018). TMEM55A overexpression does not affect lysosomal positioning (Willett et al., 2017), suggesting it may have functions independent of lysosomal positioning. In summary, the extent to which TMEM55A and TMEM55B share joint molecular functions remains unknown.

Here, we investigate the function of TMEM55B and its role in lysosomal positioning *in vivo* in neuronal cells by analyzing *Tmem55b* knockout (KO) mice, rat primary neurons, and *C. elegans* deficient for the TMEM55B/A homolog Y71H2AM.2. We found that, in contrast to non-neuronal cells where primarily retrograde transport of lysosomes is impaired upon the depletion of TMEM55B, TMEM55B depletion impaired bidirectional lysosomal transport in the mid- and distal axon, and a reduced number and abundance of lysosomes in the mid and distal axon. The conditional KO of *Jip4* resembles the phenotype resulting from TMEM55B deficiency in mice, suggesting at least partial critical dependence of LAMP1^+^-vesicles on the TMEM55B and JIP4 complex. While *Tmem55a* KO mice show no phenotype at all, double KO mice for *Tmem55b* and the close paralog *Tmem55a* show retarded growth and premature death, consistent with partially overlapping or additive functions of TMEM55A and TMEM55B. Importantly, our findings extend the knowledge of TMEM55B-dependent lysosomal positioning and reveal a function in bidirectional retrograde and, notably, anterograde transport.

## Material & Methods

### Reagents, plasmids, and antibodies

Antibodies: Analytical-grade chemicals were purchased, if not stated otherwise, from Sigma-Aldrich (MO, USA). The following antibodies were used: TMEM55B (rabbit polyclonal, 23992-1-AP, Proteintech, Rosemont, IL), LAMP1 (rat monoclonal, clone 1D4B, DSHB Hybridoma Bank, Iowa), Ankyrin G (mouse monoclonal, NeuroMAb #75-146), Actin (Sigma-Aldrich), p62 (rabbit polyclonal, BML-PW9860, Enzo, NY), GAPDH (mouse monoclonal, clone 6C5, sc-32233, Santa Cruz Biotechnology), Cathepsin D (rabbit polyclonal, in house (Jungnickel et al., 2024)), NeuN (mouse monoclonal, MAB377, clone A60, Sigma Aldrich), GFAP (mouse monoclonal, G3893, clone G-A-5, Sigma-Aldrich), Iba1 (rabbit polyclonal, GTX100042, Genetex), Ng2 (rat monoclonal, gift from Jacqueline Trotter), CD68 (rat monoclonal, clone FA-11, BioLegend, San Diego, CA), LAMP1 (mouse monoclonal, clone Ly1C6, Enzo Life Sciences Cat# ADI-VAM-EN001-D, RRID: AB_2038958, 1/250), TRIM46 (rabbit polyclonal, in-house (1/1000)).

### Mouse strains

The generation of *Tmem55a^−/−^ KO* and *Tmem55b^−/−^* KO mice (official gene names are *Pip4p1* and *Pip4p2*, respectively) is described below. *Tmem55a^−/−^*and *Tmem55b^−/−^* double KO mice were obtained by breeding the two lines. Conditional KO mice for *Jip4* (synonymously “JLP”) were described previously (Sato et al., 2015). The breeding of the *Tmem55a/b^−/−−/−^* double-KO line was approved by the “Ministerium für Landwirtschaft, ländliche Räume, Europa und Verbraucherschutz” under reference number 31-4/21. Transcardial perfusion was approved under the reference number 8-2/23V. Mice were housed under specific pathogen-free conditions (12 hours’ light/dark cycle, constant room temperature (22°C +/−2 °C) and humidity (55% rH +/− 10%)). Food and water were available ad libitum. Mice of mixed genders were used throughout the manuscript if not stated otherwise (for behavioral testing, only females were used; for body weight development, only males were used).

### Plasmids and shRNA constructs

TMEM55A-GFP and TMEM55B-GFP were cloned by insertion of the full-length murine TMEM55A / TMEM55B cDNA (Rudnik et al., 2022) into the pEGFP-N1plasmid. LAMP1-RFP was a gift from Walther Mothes (Addgene plasmid # 1817); LAMP1-GFP was a gift from Dr. Juan Bonifacino.

The following sequences for rat shRNAs inserted into pLKO.1-puro were used in this study: scrambled shRNA (5’-GCGAAACAGACTGGCATACTT-3’), shTMEM55B#1 shRNA (5’-TGCATCAGCATGTAGTCAAAT-3’), shTMEM55B#2 shRNA (5’-TCTGTGGACATTGCAAGAATA-3’). Both shRNAs have been validated using immunofluorescence in this study.

CTNS-1: (KG#371, (Edwards et al., 2013); Addgene #110882. Shortened unc-129 promoter (KG#230) was a gift from Kenneth Miller (Addgene plasmid #110925; http://n2t.net/addgene:110925; RRID: Addgene_110925). myo2p::GFP in the pBN41 plasmid (Addgene #86716).

### RNA isolation from brain hemispheres

For RNA isolation, frozen brain hemispheres were ground into a powder in liquid nitrogen (for bulk RNA-sequencing) or in a bead mill (for Nanostring nCounter analysis). The homogenized brain was used as the starting material for RNA extraction according to the instructions in the protocol for the NucleoSpin® RNA Plus Kit (Macherey-Nagel).

### Bulk RNA sequencing

Whole Transcriptome analysis was performed via RNA sequencing on a HiSeq 4000 (Illumina, San Diego, CA, USA), utilizing the totalRNA library protocol (Illumina) with 1×50 bp reads according to the manufacturer’s guidelines. Raw reads were forwarded for trimming and exclusion of low-quality reads via cutadapt (Kechin et al., 2017), followed by alignment to the mouse genome (version Ensembl Genes 92 / Mouse genes GRCm38p6) via TopHat2 (Trapnell et al., 2012). Finally, quantitative gene expression was assessed employing HTSeq (Anders et al., 2015) with subsequent quantile normalization (Bolstad et al., 2003).

### nCounter transcriptomic analysis

Total RNA was isolated from frozen brains of 14-month-old mice using the Nucleospin® RNA Midi Kit from Machery & Nagel according to the manufacturer’s protocol. 50 ng of RNA was used for gene expression profiling using the nCounter Analysis (Neuroinflammation Panel) from NanoString Technologies, Inc. The NanoString data were analyzed and normalized using nSolverTM software (version 4.0, NanoString Technologies, Inc.). RNA nCounts were normalized using the geometric mean of 10 housekeeping genes (*Aars, Ccdc127, Cnot10, Csnk2a2, Fam104a, Lars, Mtol, Tada2b, Tbp, and Xpnpep1*) using nSolverTM software.

### SDS-PAGE and immunoblot

Cell lysates were prepared as described previously (Damme et al., 2011). Proteins were separated on 10-12.5% SDS-PAGE gels and transferred to nitrocellulose membranes (Whatman, GE Healthcare, 10426994) by semi-dry blotting. The Amersham ECL Advanced Western Blotting Detection kit (GE Healthcare, RPN2135) was used for the detection of the antibody complexes.

### Generation of Tmem55a and Tmem55b knockout mice

Mice with a null mutation in the *Tmem55a* or *Tmem55b* gene were generated in a C57BL/6N background using a clustered regularly interspaced short palindromic repeats (CRISPR) genome-editing system. For this purpose, we combined Cas9 with a single-guide RNA (sgRNA) targeting the second exon of the *Tmem55a* or *Tmem55b* gene, respectively, with the following spacer sequences: sgRNA_Tmem55a: CACACACGACAGTTGATCAC; sgRNA_Tmem55b: ACGCGACAGGTAATCATGG, and microinjected them into fertilized oocytes. Genome editing was confirmed by PCR amplification in the resulting founder mice. Founder mice were screened by analyzing PCR amplicons of the target loci by native polyacrylamide gel electrophoresis (PAGE), and confirmed by Sanger sequencing. Founder mice harboring an eight-bp deletion within the second exon of *Tmem55a* and a four-bp deletion in exon two of *Tmem55b*, each leading to frameshift mutations, were selected for further breeding.

### Genotyping of Tmem55a and Tmem55b KO mice

For genotyping, tail DNA was prepared with the “DirectPCR Lysis Reagent” (Viagen Biotech). Genotyping PCRs were performed with the following primer pairs: Tmem55a_for: CCACCTCCGTATACAGCCAT; Tmem55a_rev: GCCACTTACCGTAGCTTCAT; Tmem55b_for: CCCTTGACTAGCCCGGATAG; Tmem55b_rev: CTTCGTTGCAGACACCACAT. In order to obtain a sufficient separation of the small PCR products and to detect the minor differences between the wildtype alleles and the KO alleles (eight bp for *Tmem55a* and four bp for *Tmem55b*, respectively), the PCR products were separated by capillary electrophoresis on a QIAxcel Advance System (QIAGEN, Hilden, Germany) combined with the QIAxcel DNA High-Resolution Kit (1200) from (QIAGEN, Hilden, Germany).

### Primary rat neurons

Primary neuron experiments were approved by the DEC Dutch Animal Experiments Committee (Dier Experimenten Commissie), performed in line with institutional guidelines of the University of Utrecht, and conducted in agreement with Dutch law (Wet op de Dierproeven, 1996) and European regulations (Directive 2010/63/EU). The animal protocol has been evaluated and approved by the national Centrale Commissie Dierproeven (Central Authority for Scientific Procedures on Animals; license AVD10800202216383). Female pregnant Wistar rats were obtained from Janvier, and embryos (both genders) at the embryonic (E)18 stage of development were used for primary cultures of hippocampal and cortical neurons. The animals, pregnant females and embryos, have not been involved in previous procedures.

### Primary neuron culture and transfection

Hippocampi from embryonic day 18 rat brains were dissected and dissociated with trypsin for 15 min and plated on coverslips coated with poly-L-Lysine (37.5 µg/ml) and laminin (1.25 µg/ml) at a density of 100,000/well (12-well plates). The day of neuron plating corresponds to day in vitro 0 (DIV0). Neurobasal medium (NB) supplemented with 2% B27 (GIBCO), 0.5 mM glutamine (GIBCO), 15.6 µM glutamate (Sigma), and 1% penicillin/streptomycin (GIBCO) was used to maintain neurons incubated under controlled temperature and CO_2_ (37 °C, 5% CO_2_). Hippocampal neurons were transfected at 5 DIV (for KD experiments) or 7 DIV using Lipofectamine 2000 (Invitrogen). Briefly, DNA (50 – 1000 ng/well) was mixed with 1.2 µL of Lipofectamine in 100 µL of NB; for KD experiments, 3.3 µL of Lipofectamine was used. DNA/lipofectamine mixture was incubated at room temperature for 20 min, added to neurons in NB, and incubated for 1 hour at 37 °C, 5% CO_2_. Neurons were washed three times with NB, and half of the original medium was added together with freshly prepared Neurobasal medium. Experiments were performed either within 1 day post-transfection (DIV8) or, for KD experiments, 3 days post-transfection (DIV8).

### Immunofluorescence staining and imaging of primary rat neurons

Primary neurons were incubated at room temperature with 4% paraformaldehyde supplemented with 4% sucrose in PBS for 10 min for fixation. Cells were permeabilized and blocked with 0.05% saponin supplemented with 0.2% porcine gelatin in PBS for 1 h at room temperature. Cells were incubated with primary antibodies overnight at 4 °C, washed three times with PBS, incubated with secondary antibodies for 45 min at room temperature, and washed three times with PBS. Cells were mounted in Fluoromount-G Mounting Medium (ThermoFisher Scientific). Cells were imaged using a confocal laser-scanning microscope (LSM900, with Zen (blue edition) imaging software version 3.7.97.07000 (Zeiss)) equipped with a Plan-Apochromat ×40 NA 1.30 oil DIC objective and Plan-Apochromat ×63 NA 1.40 oil DIC objective. For KD experiments, only cells displaying continuous labeling of the cytosolic fluorescent protein (fill) along the somatodendritic and axonal domains were imaged.

### Live-cell imaging of primary rat neurons

For live-cell imaging experiments, an inverted microscope Nikon Eclipse Ti-E (Nikon), equipped with a Plan Apo VC ×100 NA 1.40 oil objective (Nikon), a Yokogawa CSU-X1-A1 spinning disk confocal unit (Roper Scientific), a Photometrics Prime BSI cMOS camera (Roper Scientific), and an incubation chamber (Tokai Hit) mounted on a motorized XYZ stage (Applied Scientific Instrumentation) was used. MetaMorph (Molecular Devices) version 7.10.2.240 software was installed for controlling all devices. Coverslips mounted in a metal ring and supplemented with the original medium from neurons were imaged in an incubation chamber that maintains optimal temperature and CO_2_ (37 °C and 5% CO_2_). To visualize multiple fluorescently labeled proteins and probes, sequential imaging was used, and each laser channel was exposed for 150 – 200 ms. Neurons were imaged every 1 s for 180 s. To identify the axon, neurons were either co-transfected with cytosolic HaloTag fill and identified by morphology or incubated with a CF640R-conjugated antibody against the AIS protein neurofascin (NF-640R) for 30 min before live-cell imaging. To label HaloTag, JFX650-ligand was incubated (100 nM in NB medium) for 10 min prior to imaging, washed three times with warm NB, and returned to the original medium. The proximal axon was imaged approximately 50 µm away from the AIS, the mid axon was imaged approximately 400 to 500 µm away from the soma, and the axonal tip was traced from the mid axon and imaged at the furthest possible end.

### Image analysis and quantification of primary rat neurons

*Kymograph analysis:* Kymographs were generated from live cell images using Fiji. Segmented lines were drawn and straightened along the axon identified by morphology using a fill and/or using the AIS marker NF-CF640R. Straightened axons were re-sliced, followed by z-projection to obtain a kymograph. A random 30 μm segment was cropped for analysis. Anterograde movements were oriented from left to right in all kymographs. The recording time and segment length are indicated in each kymograph. The number of events for anterograde, retrograde, stationary, and the total number of LAMP1-GFP in scrambled and TMEM55B-depleted conditions were obtained from three independent experiments. To measure average trajectory velocity, straight-line ROIs were manually drawn along mobile segments of individual traces to obtain event length and angle for velocity calculations.

### Axonal tip LAMP1 accumulation under TMEM55B overexpression

Axons of primary neurons expressing LAMP1-RFP under GFP control or TMEM55B-GFP were first identified with the AIS marker TRIM46 and traced with a HaloTag fill to the tip. Imaging settings were kept constant across all experimental conditions. To quantify axonal tip accumulation, LAMP1-RFP intensity at the axonal tip (AT) and axon proximal to the tip (A) was measured to calculate the ratio AT/A to reflect the propensity of LAMP1 accumulation.

### Number of LAMP1 tubulation over 180s

LAMP1-GFP-expressing neurons in both scrambled and TMEM55B-depleted conditions were imaged for 180s at the soma. Lengths of elongated LAMP1 were first measured manually in Fiji; tubules that met the minimum length requirement of 1 µm were then used for quantification over 180 s.

### Immunohistochemistry

Mice were deeply anesthetized and perfused transcardially with 0.1 M phosphate buffer (PB), pH 7.4, until the liver was pale, followed by perfusion with 30 ml of 4% formaldehyde solution (4% (w/v) paraformaldehyde in 0.1 M PB). The brain and the spinal cord were removed and postfixed by immersion for 4 hours with 4% formaldehyde solution (in 0.1 M PB). Brains were cut sagittally in two halves and were stored in 0.1 M PB or directly transferred to cryoprotection solution (30% (w/v) sucrose in 0.1 M PB pH 7.4). After incubation in cryoprotection solution overnight and for at least 16 hours on a roller at 4 °C, brain hemispheres were frozen on a layer of frozen 5% sucrose on a dry-ice pre-cooled specimen mount of a Leica 9000s sliding microtome (Leica, Wetzlar, Germany). 35 µm-thick sagittal sections were prepared and collected in 0.1 M PB pH 7.4 supplemented with 0.02% Na-Azide. For the immunohistochemistry procedure, sections were transferred to 0.1 M PB, incubated in blocking solution (0.5% Triton-X 100 (v/v), 4% normal goat serum (v/v) (Gibco) in 0.1 M PB pH 7.4) for 1 ½ −2 hours and subsequently in the diluted antibody solution (antibodies in blocking solution) overnight at 4 °C. The next day, sections were washed with wash solution (0.1 M PB pH 7.4 containing 0.25% Triton-X100) three times for 15 minutes on a shaker, incubated in the secondary antibody solution (AlexaFluor, Molecular Probes/Invitrogen) for 1 ½ hours, washed three times again for 15 minutes in the wash solution, and finally washed in 0.1 M PB pH 7.4 for ten minutes (all steps at room temperature). The slides were transferred to glass slides with a brush, dried, and embedded in Mowiol/DABCO containing 4’,6-Diamidino-2-phenylindole (DAPI). Sections were imaged using a Zeiss confocal microscope (Zeiss LSM980 AiryScan 2) equipped with a 63x objective and an automated stage. ZEN 3.3 Blue Edition software (Zeiss) was used to process the images.

Spinal cord slices were imaged using a slide scanner (Zeiss Axioscan 7) equipped with a 20x objective and automatically stitched. Spinal cord slices were further imaged using a Leica confocal microscope (Leica Stellaris 8 Falcon) equipped with a 40x objective for higher resolution. Images were processed with the ImageJ Fiji.

### Determination of the activity of lysosomal enzymes

The *in vitro* activity of the lysosomal hydrolases α-mannosidase, β-galactosidase, β-glucuronidase, and β-hexosaminidase was measured in cell or organ lysates using the artificial and chromogenic substrates p-nitrophenyl-α-D-mannopyranoside, p-nitrophenyl-β-D-galactopyranoside, p-nitrophenyl-β-D-glucuronide, and 4-nitrophenyl-N-acetyl-β-D-glucosamine, respectively. Enzyme-mediated hydrolysis of the substrates leads to the formation of 4-nitrophenol, which can be detected and quantified photometrically at 405 nm under alkaline conditions.

The measurement was performed in duplicate, with 10 μL each of brain lysate (protein concentration ∼10 μg/μL, prepared as described for SDS-PAGE) in 0.2 M trisodium citrate, pH 4.6, 0.4% (w/v) BSA, and 0.08% (w/v) NaN_3_ for 5 min to 6 h at 37 °C. The reaction was stopped by adding 500–1000 μL of 0.4 M glycine solution (pH 10.4), and the insoluble aggregates were pelleted at 16,000×g at room temperature for 10 min. The supernatants were used for photometric measurement of absorbance at 405 nm using a Synergy HAT (BioTek). Each lysate sample was also incubated and measured without the corresponding substrate as a negative control/substrate blank.

### Behavioral testing of mice

All behavioral tests were performed under standardized conditions with female wildtype and *Tmem55b^−/−^*mice. Behavioral testing was approved by the Ethical Review Board (ECD) of the University of Leuven that works in accordance with European Directive 2010/63/EU.

### Forelimb grip

Forelimb grip strength was evaluated by allowing mice to spontaneously grab a T-shaped bar. Connection of the bar to a digital dynamometer (Ugo Basile, Gemonio, Italy) allowed quantification of strength, which was averaged over 10 trials.

### Rotarod test

The rotarod test measured motor coordination and equilibrium. Animals first received 2 minutes of training at a fixed speed of 5 rpm on the apparatus (Ugo Basile, Gemonio, Italy). Following training, 4 test trials (10 min intertrial interval) were conducted with an accelerating rotation from 5 to 40 rpm during 5 minutes. Drop latency was registered up to the 5 min cut-off.

### Vertical pole test

Motor function was further evaluated using the vertical pole test. Mice were positioned with their head upwards on the top side of a vertical wooden beam, the surface of which was made rough with rope (length 55 cm, cross-section 1.5 cm). Latency to turn downward and descend was registered for every animal.

### Footprint gait analysis

Footprint gait analysis was performed via ventral plane video-tracking during treadmill walking for automatic quantification of gait patterns (Digigait, MouseSpecifics Inc, MA, USA). Mice were habituated to walking on the treadmill at 10 cm/s. Belt speed was gradually increased to 20 cm/s from the moment video data were acquired for 45 s. The number of times mice could not keep pace with the treadmill speed was registered. A section of the video in which the mice consistently walked above the camera was selected for analysis. Different parameters were extracted from the video data using Digigait analysis software, including fore vs hind gait symmetry (the ratio of forelimb stepping frequency to hind limb stepping frequency). Spatial parameters comprised stance widths (distance between contralateral paws), stride lengths (distance between subsequent placements of the same paw), and step angles (angles between left and right limbs as a function of stance width and stride length). Temporal aspects of gait were also evaluated by analyzing stance (time paw is in contact with the belt), brake (part of stance from initial contact to maximum contact), propulsion (part of stance from maximum contact to swing), swing (time paw is not in contact with the belt), and stride (sum of stance and swing duration) times.

### Balance beam test

The balance beam test was performed to gain further insight into sensorimotor function. Mice were trained to walk across a set of 1 m long narrow beams. Six beams were used, whose shape and diameter represented increasing challenge for balance and equilibrium (square (SQ), cross-section: 28, 12 & 5 mm; round (RO), diameter: 28, 17 & 11 mm). Beams were placed 50 cm above the surface, terminating on a square escape platform. Mice underwent a training phase during which they learned to traverse the 12 mm square beam without hesitation. The test phase consisted of 2 consecutive trials on each beam in the order described above. Latency to cross and number of hind paw slips were recorded for each trial. The average of both trials was used for analysis.

### Home cage activity

Home cage activity was recorded in 20 x 30 cm transparent cages, placed between 3 infrared beams. Activity patterns were evaluated by recording the total number of beam crossings during 30 min intervals for 23 h using customized software.

### Open field test

Open field locomotion was recorded to evaluate exploratory behavior. Following 30 min dark adaptation, mice were placed into a transparent 50 × 50 x 30 cm arena. Subsequent to 1 min habituation, animal movement was recorded for 10 min using ANY-mazeTM Video Tracking System software (Stoelting Co., IL, USA). Total path length, time in center (= 30 cm circle), and time in the periphery (< 5 cm from the walls) were extracted.

### Elevated plus maze test

Exploration of an elevated plus maze was assessed to characterize anxiety-related behavior. The arena consisted of a plus-shaped maze, elevated 50 cm above the surface, with two arms (5 cm wide) closed by side walls and two open arms. Mice could freely explore for 10 minutes following 1 min habituation. Four IR beams recorded open and closed arm entries, and one recorded the percentage of time per min spent in the open arms, and were connected to a computerized activity logger.

### Sociability and preference for social novelty

Sociability and preference for social novelty (SPSN) were evaluated in a three-chamber Plexiglas set-up. The two side chambers (10 x 10 cm) were separated from the center compartment (40 x 10 cm) by Plexiglas walls containing holes (diameter 0.8 cm) to allow social interaction and odor transfer between experimental and stimulus mice. First, an acclimation trial was performed during which mice were allowed to explore the central chamber for 5 min. During the 10 min sociability trial, one side chamber contained a stranger mouse. The side of the social chamber was randomized. Preference for social novelty was subsequently evaluated by placing a second stranger mouse into the previously empty side chamber for 10 min. Animal movement was recorded using ANY-mazeTM Video Tracking System software (Stoelting Co., IL, USA). The time the animal’s head was closer than 1 cm to the chamber walls was measured to allow quantification of preferential exploration in sociability and social novelty phases.

### Novel object recognition test

The novel object recognition test evaluated the ability of mice to recognize new objects in their environment. During the first day, mice could freely explore a transparent 50 × 50 x 30 cm arena for 10 min (habituation phase). During the second day, mice were placed for 10 min in the same arena, which now contained 2 identical objects (X-X) placed near opposite corners of the set-up (familiarization phase). After approximately 3.5 hours, mice were again placed in the arena for 10 min with 2 objects. However, 1 object was identical to the objects from the familiarization phase, and 1 object was a different, unfamiliar object (novel object phase). Objects were filled in 50 ml Falcon tubes and similar-sized towers of Duplo blocks. The location of the novel and familiar object was counterbalanced across the experimental groups. Animal movement was recorded using ANY-mazeTM Video Tracking System software (Stoelting Co., IL, USA). The time the head of the animal was within the predefined object zone (10 x 10 cm) was recorded to quantify familiar vs novel object exploration.

### Passive avoidance learning test

Passive avoidance learning was examined in a step-through box with a small illuminated compartment and a larger dark compartment containing a grid floor. After 30 min dark adaptation, the mouse was placed in the light compartment for a training trial. After 5 s, the sliding door to the dark compartment was opened, and step-through latency was manually recorded up to a 300-s cut-off. When all four paws were placed on the grid, a mild electric shock (0.3 mA, 1 s) was delivered with a constant current shocker (MED Associates Inc., St. Albans, Vermont, USA). Retention memory was tested 24 h later using the same procedure without shock delivery.

### Tail withdrawal test

The tail withdrawal test was performed to evaluate the response to nociceptive stimuli. Mice were placed in a custom-made restrainer, which allowed the distal half of the tail to be placed in warm water (50-52 °C). Latency to respond to the heat stimulus with vigorous flexion of the tail was measured.

### Caenorhabditis elegans strains and culturing

All *C. elegans* strains were grown and fed ad libitum at 20 °C on nematode growth medium (NGM) agar plates seeded with *Escherichia coli* OP50 as a food source (Brenner, 1974). The Bristol N2 (used as wildtype) was provided by the *Caenorhabditis Genetics Center* (CGC). The *Y71H2AM.2* deletion mutant allele (syb1124) was generated by SunyBiotech Ltd. using CRISPR/Cas9-mediated deletion of the entire *Y71H2AM.2* genomic sequence in the N2 wildtype background. *Y71H2AM.2(syb1124)* mutant hermaphrodites were subsequently outcrossed to N2 males four times. Homozygosity of the mutant allele was confirmed by PCR amplification of genomic DNA and/or sequencing (Eurofins Genomics GmbH) of the resulting PCR products. PCR proof of the deletion was performed using two different primer. One primer pair was designed to flank the *Y71H2AM.2* genomic sequence at its 5’ end (GCATTGTGGTTACCATAGGCAT) and 3’ end (TAGAAAGTTGTCCGAAATCGCG). The second pair was designed with one primer positioned within the genomic sequence (exon 3; caactgttgctgtgttggctg) and the other outside the 3’ end (aactcgaacccactcaactcc). For clarity, the *Y71H2AM.2(syb1124)* deletion allele is referred to as *Y71H2AM.2(null)* throughout this study.

### Molecular biology and transfection of C. elegans

Molecular cloning was performed using standard PCR-based cloning procedures according to the Addgene protocol and as previously described (Edwards et al., 2015). For lysosomal localization and colocalization experiments in motorneurons, an expression construct encoding the lysosomal marker cystinosin (CTNS-1; KG#371; Addgene) fused to a C-terminal mCherry reporter (1.97 kb) and the *unc-54* 3′ UTR (0.9 kb) was expressed under the control of the motor neuron-specific *unc-129* promoter (2.64 kb) in N2 wildtype and *Y71H2AM.2(null)* animals (*unc-129p::ctns-1::mCherry::unc-54* 3′ UTR).

For neuron-specific rescue experiments, extrachromosomal constructs were generated using the KG#230 plasmid backbone (Addgene). The *C. elegans Y71H2AM.2* genomic sequence or its *Mus musculus* ortholog *Tmem55b* was inserted into the multiple cloning site using BamHI and NheI restriction sites. Each sequence was placed downstream of the motor-neuron-specific *unc-129* promoter (2.64 kb; PsiI/BamHI) and fused at its 3′ end to GFP (0.62 kb; NheI/SalI), followed by the *unc-54* 3′ UTR (0.3 kb; SalI/SpeI). Restriction sites required for cloning were introduced by PCR using primers containing the corresponding restriction site sequences.

The neuron-specific rescue of the Y71H2AM.2-deficient worms was performed *via* genetic constructs based on the KG#230 plasmid backbone *(Addgene)* containing the *C. elegans* genomic sequence of *Y71H2AM.2* or its *Mus musculus* ortholog TMEM55B, both inserted in the multiple cloning site using the *BamHI* and *NheI* restriction sites. Both sequences are flanked at 5’ by the *unc-129* (2.64 kb fragment) promoter (restriction sites: PsiI, BamHI) and fused at 3’ with a C-terminal GFP (0.62 kb; NheI, SalI) and an *unc-54* 3’UTR (0.3 kb; restriction sites: SalI, SpeI). The restriction sites of the cloned genomic sequences were integrated by PCR with primer pairs designed to contain the corresponding restriction sites.

Transgenic strains were generated by biolistic bombardment as previously described (Wilm et al., 1999). For rescue experiments, *myo-2p*::GFP from the pBN41 plasmid (Addgene) was used as a co-injection marker. For all constructs generated from PCR-amplified fragments, the integrity of the cloned sequence was verified, and only clones containing no mutations within the sequence of interest were used.

### Imaging of C. elegans reporter strains

Lysosomal distribution and migration phenotypes were assessed *in vivo* using a Zeiss Axio Imager Z2 upright microscope equipped with an LSM 700 laser-scanning confocal system (Carl Zeiss, Germany). GFP and mCherry fluorescence was excited at 488 and 555 nm, respectively. For imaging, groups of 10 age-synchronized transgenic animals were transferred onto glass slides containing 2% agarose pads and immobilized with 2 μl of 5 mM levamisole. Samples were covered with a coverslip and immediately mounted on the microscope for image acquisition. Confocal images were acquired and processed using ZEN software (Carl Zeiss, Germany). Identical imaging parameters were used across experimental groups within each experiment to enable direct comparison of fluorescence patterns.

For time-lapse imaging of CTNS-1::mCherry particles in the DA9 axon, L4 hermaphrodites were paralyzed with 5 mM levamisole in M9 buffer for 10 min, transferred to a freshly prepared 2 % (w/v) agarose pad, and covered with an 18 mm x 18 mm coverslip (No. 1.5H, Marienfeld). Time-lapse sequences of the axon initial segment were recorded at 20 °C on a Nikon Eclipse Ti microscope coupled to an Andor Revolution XD spinning disk confocal system, composed of an iXon Ultra 897 CCD camera (Andor Technology), a solid-state laser combiner (ALC-UVP 350i, Andor Technology), and a CSU-X1 confocal scanner (Yokogawa Electric Corporation). The system was controlled by Andor IQ3 software (Andor Technology). A single image was acquired every 500 ms for a total of 90 s at 1 x 1 binning using a 100x NA 1.45 Plan-Apochromat objective. Kymograph generation and analysis were performed with Fiji software (ImageJ version 1.52d) with the KymoAnalyzer macro package.

### C. elegans locomotion assays and analyses

Age-dependent locomotor activity on nematode growth medium (NGM) agar plates was assessed as previously described (Gottschling et al., 2017; Luersen et al., 2016), with modifications. For all locomotion assays, mutant and transgenic strains were analyzed in parallel with their respective control strains (N2 wildtype animals or non-transgenic siblings) on the same day and using plates prepared from the same batch and seeded with *Escherichia coli* OP50.

To assess age-dependent locomotor activity, age-synchronized, well-fed animals were analyzed at days 3, 10, and 15 after synchronization by bleaching. Individual animals were transferred in 10 μl of M9 buffer onto locomotion assay plates seeded with a thin lawn of *E. coli* OP50. Plates were seeded with 100 μl of an overnight OP50 culture per 60-mm plate and incubated for 12 h at 37°C before use. Animals were allowed to move freely on the assay plates for 1 h at 20 °C. Locomotion tracks from at least 10 individual animals per condition were assessed in each experiment, and at least three independent experiments were performed.

For track analysis, locomotion traces were marked on the underside of each NGM plate and subsequently imaged using a VRmagic C-9+/BW PRO IR-CUT camera (VRmagic, Germany) mounted on a Zeiss Stemi 2000-C stereomicroscope (Carl Zeiss, Germany). Images were saved as JPEG files, and locomotion tracks were ranked by visual inspection. Track lengths were quantified using ImageJ.

### C. elegans life-span assays and analyses

Lifespan assays were performed as previously (Kenyon et al., 1993). Animals were age-synchronized and maintained at 20°C on NGM agar plates seeded with *E. coli* OP50 (Kenyon, 1988; Stiernagle, 2006). Mutant strains and N2 wildtype controls were assayed in parallel to minimize potential effects of environmental variation. For each assay, 10 *ad libitum*-fed L4-stage animals were transferred to 90-mm NGM plates to minimize overcrowding and competition for food during adulthood. Animals were scored every 24 h as alive, dead, or lost. During the reproductive period, animals were transferred daily to freshly seeded plates and subsequently transferred every 72 h. Animals that failed to move spontaneously or in response to gentle stimulation with a platinum wire were scored as dead. Animals that were lost during the experiment were censored from the analysis. At least three independent experiments were performed for each strain.

### Statistical analyses

Statistical analyses were performed using GraphPad Prism version 9.03. The statistical tests used for individual experiments are specified in the corresponding figure legends. Unless otherwise indicated, comparisons between two groups were performed using an unpaired two-tailed Student’s *t*-test, whereas comparisons among multiple groups were performed using ordinary one-way ANOVA. Data are presented as mean ± SEM unless otherwise indicated. Statistical significance was defined as *p* < 0.05. Significance levels are indicated as follows *, p < 0.05; **, p < 0.01; ***, p < 0.001; ****, p < 0.0001.

## Results

### TMEM55B is highly expressed in neurons of the central nervous system

TMEM55B plays a critical role in the microtubule-dependent retrograde positioning of lysosomes upon overexpression and knockdown (KD) in non-neuronal cell-based models (Willett et al., 2017). To understand the physiological function of TMEM55B *in vivo*, we first analyzed expression levels in tissues of adult mice using a TMEM55B-specific antibody (**Supplemental Figure 1A**). TMEM55B is expressed in different organs, with the highest protein levels among the analyzed tissues in the testis, the brain, followed by the spleen, the lung, and the thymus, and very low relative expression in the kidney and the liver (**Supplemental Figure 1A**). Given the high expression in the brain and the importance of lysosomal trafficking in highly complex, polarized cells such as neurons, we focused our subsequent analysis of TMEM55B function on the central nervous system. Immunofluorescence studies with a TMEM55B-specific antibody showed robust staining in the cerebral cortex, with a distribution of TMEM55B signals typical for cortical neurons (**Supplemental Figure 1B**). Co-immunofluorescence studies with cell-type-specific markers for neurons (NeuN), astrocytes (GFAP), microglia (Iba1), and oligodendrocyte progenitor cells (OPCs) (Ng2) revealed the highest expression of TMEM55B in neurons (**Supplemental Figure 1C**), compatible with their high demand for proteins related to organelle trafficking in neurites. Furthermore, a co-immunofluorescence experiment of TMEM55B and the lysosomal marker LAMP1 revealed prominent co-localization in cortical neurons *in situ*, confirming predominantly lysosomal localization at endogenous levels (**Supplemental Figure 1D**).

### CRISPR/Cas9-mediated Tmem55b^−/−^ knockout mice are viable and show altered distribution of lysosomes in the brain in situ

To study the physiological function of TMEM55B *in vivo* and dissect the physiological and intracellular processes it is involved in, we generated *Tmem55b* KO mice (*Tmem55b^−/−^*) by CRISPR/Cas9-mediated gene editing. Analysis of founder mice revealed different short Indel mutations in the targeted coding exon 2, and we selected a founder line with a four-base-pair deletion that causes a frameshift (**Supplemental Figure 2A-E**). In contrast to a previous report, in which early embryonic lethality was reported upon *Tmem55b* gene disruption (Hashimoto et al., 2018), CRISPR/Cas9-mediated *Tmem55b^−/−^* KO mice in a C57/Bl6N background were viable/born at the expected Mendelian ratios and survived to adulthood without increased mortality or any apparent gross morphological abnormalities (**Supplemental Figure 2F**). *Tmem55b^−/−^* KO mice had a lifespan comparable to wildtype littermates. The full KO was validated at the protein level by analyzing tissue lysates from the brain and different organs by immunoblot (**Figure 1A, Supplemental Figure 2G**) and by immunohistochemistry on brain sections (**Figure 1B**) with a TMEM55B-specific antibody.

**Figure 1.**
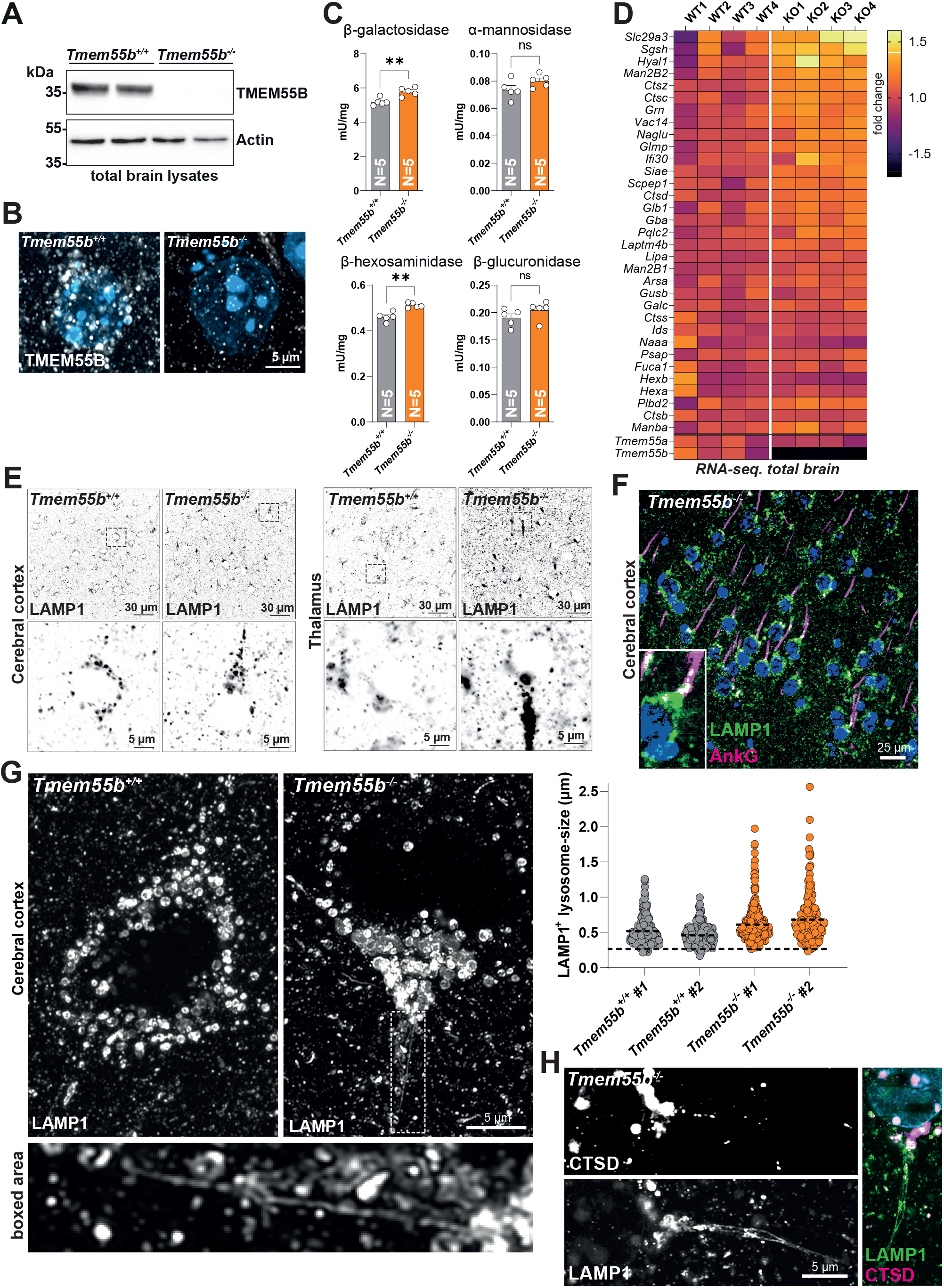
Tmem55b knockout mice accumulate enlarged and tubular LAMP1^+^-lysosomes in the PAEZ and AIS. **(A)** Immunoblot of total brain lysates from wildtype (*Tmem55b^+/+^*) and *Tmem55b^−/−^* KO (*Tmem55b^−/−^*) mice with an antibody against TMEM55B. Actin is shown as a loading control. **(B)** Immunofluorescence staining of brain sections from *Tmem55b^+/+^* and *Tmem55b^−/−^* KO mice with an antibody against TMEM55B. Nuclei are stained with DAPI (blue). A cortical neuron is shown. **(C)** The specific activity of β-galactosidase, α-mannosidase, β-hexosaminidase, and β-glucuronidase in total brain lysates (N = 5, unpaired t-test; ns = not significant, ** = p ≤ 0.01, age: 10 months). **(D)** Heatmap representing the fold changes in the transcript abundance of selected lysosomal genes/transcripts assessed by bulk RNA sequencing of total brain RNA. Age: 10 months. **(E)** Immunofluorescence staining of brain sections from *Tmem55b^+/+^* and *Tmem55b^−/−^* mice with an antibody against LAMP1. The cerebral cortex and the thalamus are shown. Age: 10 months. **(F)** Immunofluorescence staining of a brain section from a *Tmem55b^−/−^* mouse with an antibody against LAMP1 (green) and Ankyrin G (AnkG, magenta). Age: 10 months. **(G)** High-resolution Airyscan microscopy images of LAMP1 immunofluorescence staining of cortical neurons from *Tmem55b^+/+^* and *Tmem55b^−/−^* KO mice. The boxed area with a long tubular LAMP1^+^-structure is magnified. Quantification of individual lysosome size is shown; two animals per genotype were analyzed; n=218-230 organelles per animal, age 10 months. **(H)** Airyscan high-resolution microscopy of a neuron from a *Tmem55b^−/−^* mouse with an antibody against LAMP1 (green) and cathepsin D (CTSD, magenta). Age: 10 months.

In order to characterize the consequences of a loss of TMEM55B on the endo-/lysosomal system *in vivo*, we determined the specific activity of the lysosomal hydrolytic enzymes β-galactosidase, β-glucuronidase, α-mannosidase, and β-hexosaminidase in brain lysates from 10-month-old wildtype and *Tmem55b^−/−^* KO mice (**Figure 1C**). We observed minor but statistically significant differences in the specific activities of the lysosomal enzymes β-hexosaminidase and β-galactosidase, with a modest increase in brain lysates from *Tmem55b^−/−^*KO mice. Transcriptome analysis of whole-brain RNA from adult wildtype and *Tmem55b^−/−^* KO mice by bulk RNA sequencing revealed a clear separation of both genotype groups by principal component analysis (**Supplemental Figure 3A**) and revealed subtle transcriptional upregulation of a number of lysosomal genes (**Figure 1D**), consistent with modestly altered enzyme activity levels, suggesting that TMEM55B deficiency affects the endo-/lysosomal system. The transcription of *Tmem55a* was not changed, e.g., as a compensatory mechanism.

Next, to analyze the spatial distribution of lysosomes in TMEM55B-deficient neurons, we performed immunofluorescence analyses on brain sections from *Tmem55b^−/−^* KO mice for lysosomal marker proteins (**Figure 1E)**. In contrast to cortical neurons from wildtype mice, where LAMP1^+^ lysosomes were confined to the soma and distributed around the nucleus, lysosomes prominently clustered in a Y-like fashion towards the axon hillock, a region that functionally corresponds to the PAEZ (**Figure 1E)**. Co-immunofluorescence of LAMP1 with Ankyrin G confirmed the localization of the clustered LAMP1^+^-lysosomes proximal to the AIS and partially within the AIS (**Figure 1F**). The observation of clustered lysosomes was not restricted to cortical neurons but was also observed in other neuronal populations, such as hippocampal neurons, and was most pronounced in thalamic neurons (**Figure 1E)**. Analysis of animals ranging from 6 weeks to 1.5 years old revealed that the redistribution was age-independent and was an early phenotypic feature. High-resolution Airyscan microscopy of cortical sections revealed an increase in the size of LAMP1^+^ lysosomes in cortical neurons from *Tmem55b^−/−^* KO mice of 0.6 - 0.7 µm compared to 0.48 - 0.5 µm, with enlarged lysosomes >1.5 µm only seen in neurons of *Tmem55b^−/−^* KO mice. Importantly, we regularly observed conspicuous tubular, elongated LAMP1^+^ organelles extending into the AIS (**Figure 1G**). Such tubular structures were absent from cortical neurons of wildtype mice. Co-immunofluorescence of LAMP1 with the lysosomal hydrolase Cathepsin D (CTSD) revealed the presence of this soluble enzyme in a subpopulation of LAMP1^+^ organelles, and remarkably, the tubular structures were devoid of CTSD immunoreactivity (**Figure 1H**).

Confocal microscopy of unstained brain sections revealed increased abundance of autofluorescent lipofuscin in neurons from *Tmem55b^−/−^* KO mice (as shown for the hippocampus and cerebral cortex; **Figure 2A**), thereby indirectly implicating impaired endolysosomal degradation and lysosomal dysfunction. Immunohistochemical detection of p62, an autophagy cargo adaptor, revealed the presence of large (>5 µm) p62-positive aggregates in distinct layers of the cerebral cortex of aged *Tmem55b^−/−^* KO mice (**Figure 2B**), suggesting impaired turnover of such aggregates via the endo-/lysosomal system and autophagy. Interestingly, p62 often decorated LAMP1-positive vesicles, suggesting damaged lysosomes marked for autophagic turnover (lysophagy) (**Figure 2C**). While p62-aggregates were abundant in a distinct layer of the cortex, major parts of the brain lacked p62 pathology. Next, we analyzed LAMP1 levels and cathepsin D proteolytic processing by immunoblot in brain lysates as a readout of delivery of a prototypical lysosomal hydrolase (**Supplemental Figure 3B**). Total LAMP1 levels were not affected by TMEM55B deficiency, as assessed by immunoblot. Likewise, the proteolytic processing of cathepsin D was unaltered, suggesting that the delivery or maturation of soluble lysosomal enzymes is unaffected.

**Figure 2.**
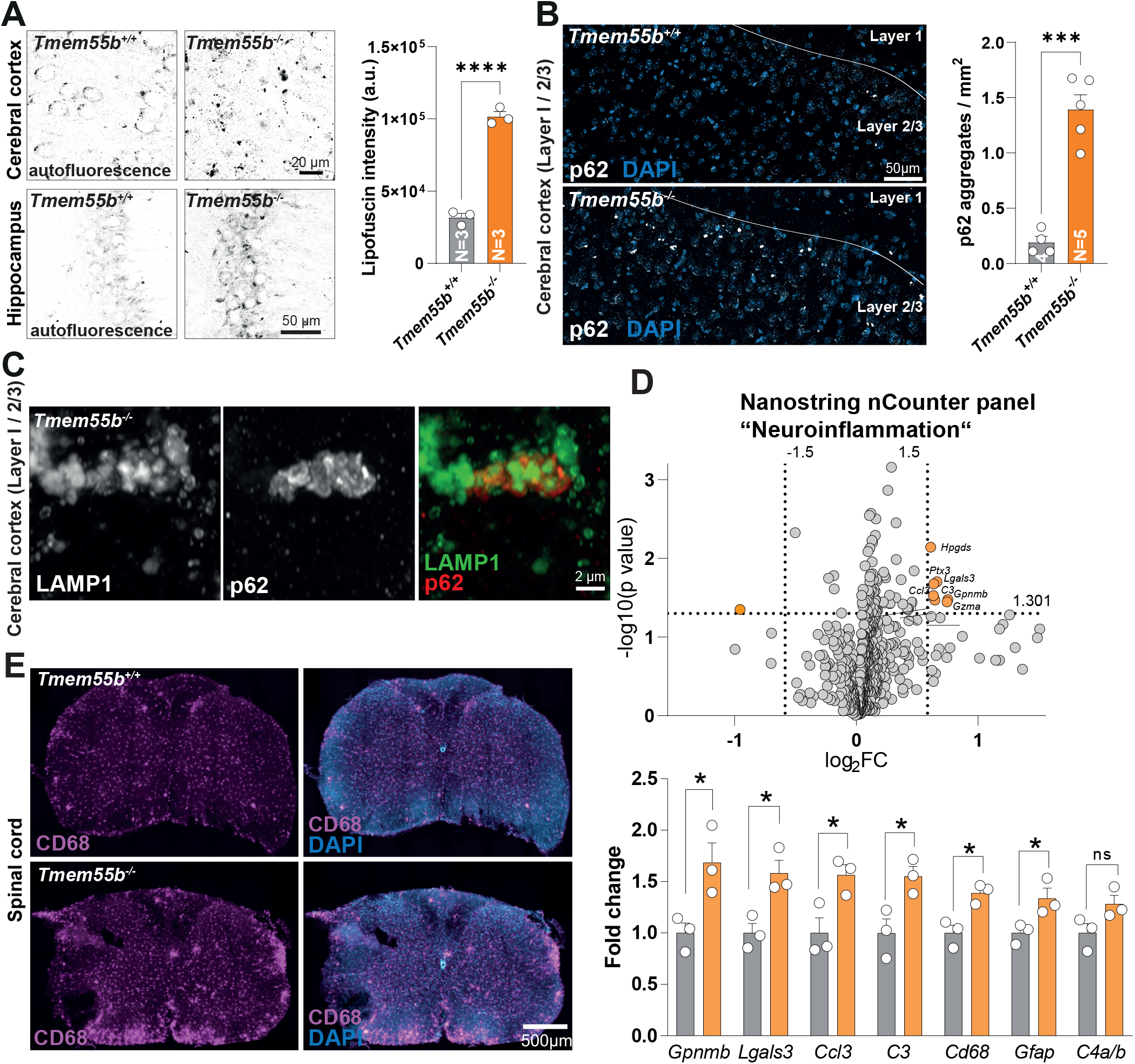
Aged *Tmem55b^−/−^* mice show lipofuscin deposition, p62 aggregates, and mild white-matter microgliosis. **(A)** Representative confocal microscopy images of unstained brain sections of the hippocampus and the cerebral cortex areas using a 488 nm laser for the detection of autofluorescent lipofuscin (Age: 14 months). Quantification is depicted for the cerebral cortex (N=3, n = 40 neurons/animal; Unpaired t-test, **** = P ≤ 0.0001). **(B)** Immunofluorescence staining of the cerebral cortex of *Tmem55b^+/+^* and *Tmem55b^−/−^* KO mice for p62 (white). Nuclei (blue) are stained with DAPI. A quantification of the number of p62-positive aggregates is shown. (N=4-5, Unpaired t-test, *** = P ≤ 0.001. Age: 24 months; Z-stack projections). **(C)** Double co-immunofluorescence staining of the cerebral cortex of *Tmem55b^−/−^* KO mice for p62 (red) and LAMP1 (green). Age: 24 months; Z-stack projections. **(D)** Volcano plot presentation of the differentially expressed transcripts of wildtype and *Tmem55b^−/−^* KO mice (N=3 / genotype) from RNA isolated from 14-month-old mice assessed by nCounter expression profiling (“Neuroinflammation”-panel). The thresholds for transcripts reaching statistically significant differences (fold change (fc) < − 1.5 / > 1.5; p < 0.05, no adjustment for multiple testing) are indicated (dotted lines), and the quadrants with transcripts reaching the threshold are colored in orange. Bar graphs are depicted for selected transcripts. **(F)** Immunofluorescence staining of spinal cord sections from wildtype and *Tmem55b^−/−^* KO mice for CD68 (magenta). Nuclei (blue) are stained with DAPI. Age: 14 months.

Lysosomes mediate the final degradation of cargo delivered by axonal autophagosomes that are retrogradely transported towards the soma upon fusion with anterogradely transported lysosomes and endolysosomes (Cason et al., 2022). We therefore analyzed steady-state LC3 levels as a readout of autophagosome number (**Supplemental Figure 3B**). Surprisingly, we did not observe the accumulation of LC3-II, the lipidated autophagosome-bound form of LC3, in brain lysates, even in aged animals (16 months), suggesting that, despite the apparent pre-axonal lysosomal positioning defects, the overall degradation of autophagosomes is not impaired in *Tmem55b^−/−^*KO mice.

To determine whether the lysosomal changes in neurons also affect glial cells - and microglia in particular (e.g., as an indication of subtle degenerative changes resulting in neuroinflammation) - we analyzed whole-brain RNA using sensitive nCounter transcriptome analysis with the “Neuroinflammation” panel (age: 14 months) (**Figure 2D**). Only a few transcripts showed fold changes >1.5-fold; these included a typical activated microglial signature, with *Gpnmb, Lgals3, Ccl3, C3*, and *Cd68* as differentially expressed transcripts. To validate these findings and add spatial context, we performed histological analysis using CD68 as a sensitive marker of microglial activation.

Immunohistochemistry of the spinal cord of 14-month-old mice revealed an increase in CD68 reactivity in the white matter of the cervical spinal cord, which was most pronounced around the ventral medial fissure (**Figure 2E**). In summary, our analyses revealed only minor and late-onset impairment of autophago-/lysosomal degradation and mild signs of microgliosis in white matter tracts, suggesting axonal insult and damage.

To analyze the consequences of a loss of TMEM55B on neuronal function, we tested adult (age: 8 months) *Tmem55b^−/−^* KO mice in a battery of behavioral tests (**Supplemental Figure 4**). No statistically significant differences compared with age-matched wildtype mice could be determined in grip strength, rotarod performance, gait symmetry, or spatial gait parameters (**Supplemental Figure 4A-C**). No significant differences were observed in the open field exploration patterns (**Supplemental Figure 4D**). *Tmem55b^−/−^*KO mice showed no deficits in the balance beam test, used to assess motor coordination and balance (**Supplemental Figure 4E**). To assess general locomotor activity and circadian activity, cage activity was determined over 23 h (**Supplemental Figure 4F**). *Tmem55b^−/−^* KO mice showed a time-dependent increase in activity, as they were more active during the first nocturnal peak (between 20 h and 23.30 h) compared to wildtype mice. There was no difference between *Tmem55B^+/+^* and *Tmem55b^−/−^* KO mice for latency to tail withdrawal, suggesting similar responsivity to a nociceptive stimulus. (**Supplemental Figure 4G**). No significant differences were observed in the elevated plus maze test (**Supplemental Figure 4H**). In summary, these data indicate that the mislocalization of lysosomes has no major influence on neuronal function and only mildly alters behavior.

### Conditional JIP4 knockout mice resemble the neuronal lysosomal positioning phenotype of Tmem55b^−/−^ KO mice

TMEM55B recruits the cytosolic scaffold protein JIP4 (Willett et al., 2017), which serves as a platform for both microtubule motor proteins dynein and kinesin-1. Interestingly, JIP4 shows a similar tissue expression to TMEM55B with robust expression in the brain (**Supplemental Figure 5A**). For a direct comparison of the depletion of TMEM55B and JIP4 in a neuron-specific background, we analyzed brain tissue from forebrain conditional *Jip4* KO mice (*Jip4^fl/fl^*EMX cre+, *Jip4* cKO), in which *Jip4* is deleted in EMX-cre-expressing forebrain neurons (Sato et al., 2015) (**Figure 3A**). Notably, the conditional *Jip4* cKO mice exhibited a strikingly similar mislocalization of LAMP1+ lysosomes comparable to *Tmem55b^−/−^*KO mice in neurons of the cortex and hippocampus, while lysosomes were normally distributed in the thalamus (**Supplemental Figure 5B**), in which the EMX promoter is not active, and thus JIP4 is not depleted. Moreover, *Jip4* cKO mice accumulated lipofuscin comparable to *Tmem55b^−/−^* KO mice. Membrane fractionation of brain homogenates by ultracentrifugation to separate membrane and membrane-associated proteins from soluble proteins of wildtype and *Tmem55b^−/−^* KO mice revealed only a minor fraction of JIP4 associated with membranes and no differences between wildtype and *Tmem55b^−/−^* KO mice (**Figure 3B**), suggesting that only a minor fraction of JIP4 associates with TMEM55B. Of note, bulk membrane fractionation does not resolve lysosome-specific pools; a lysosome-specific approach (e.g., Lyso-IP) would be required to determine whether TMEM55B specifically affects JIP4 recruitment to lysosomal membranes. In summary, these data support the conclusion that TMEM55B and JIP4 act in a shared pathway regulating lysosomal positioning in forebrain neurons, but that JIP4 fulfills other functions independent of TMEM55B-mediated lysosomal positioning.

**Figure 3.**
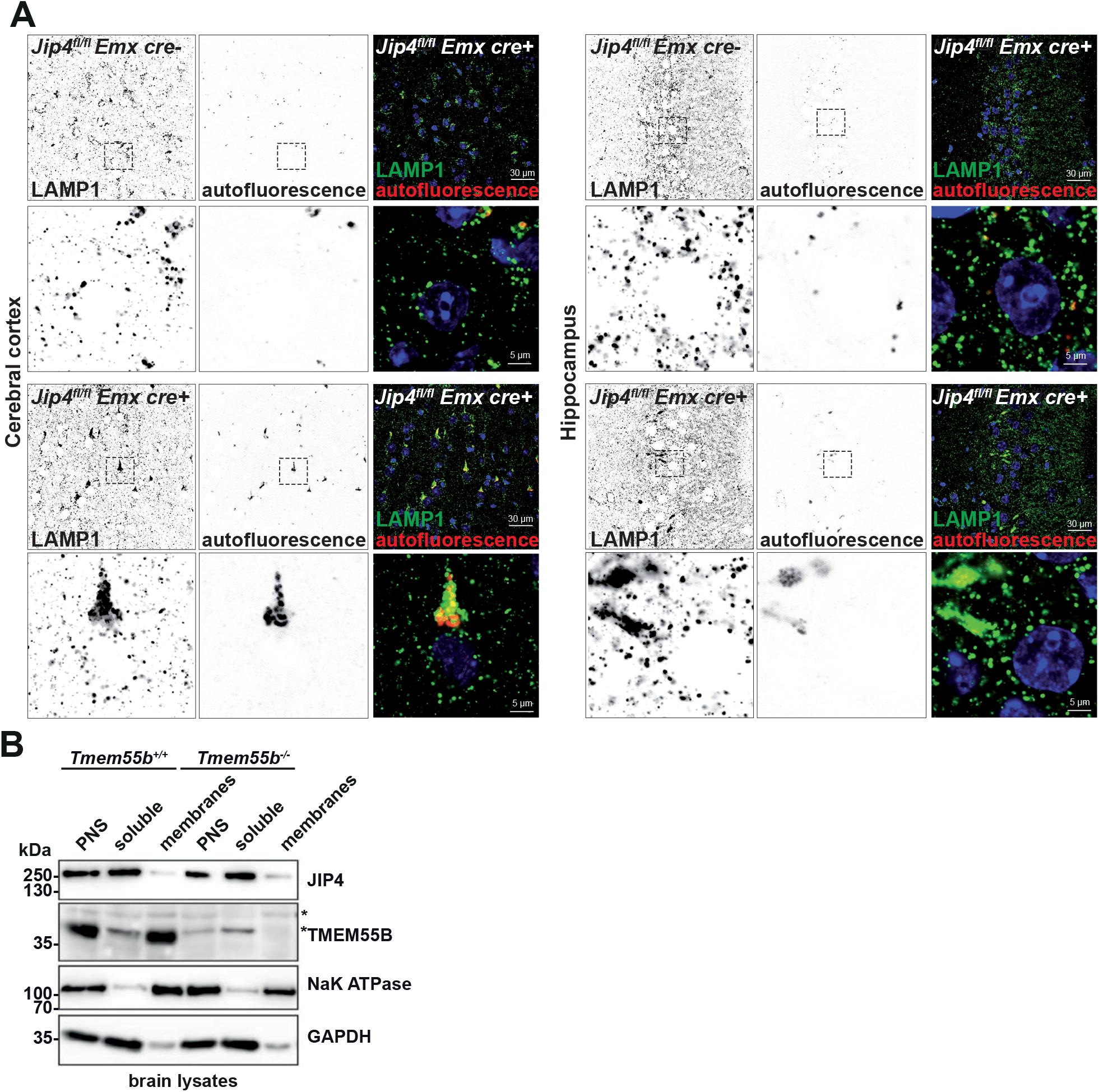
Forebrain conditional JIP4 KO mice resemble the neuronal lysosomal positioning defect of Tmem55b^−/−^ KO mice. **(A)** Immunofluorescence staining of brain sections of floxed EMX cre-negative *Jip4* mice (*JIP4^fl/fl^* EMX cre-) and EMX cre-positive conditional forebrain-specific *Jip4* KO mice (*JIP4^fl/fl^* EMX cre+) for LAMP1 (green) and autofluorescence (red) of the cerebral cortex (left) and the hippocampus (right). Age: 9 months. **(B)** Immunoblot analysis of membrane fractions of brain homogenates from wildtype and *Tmem55b^−/−^* KO mice by ultracentrifugation with antibodies against JIP4 and TMEM55B. NaK ATPase and GAPDH are shown as markers for membrane- and soluble proteins, respectively. PNS = Postnuclear supernatant.

### The dynamic distribution of lysosomes in the axon is impaired in TMEM55B-depleted primary rat neurons

Next, we analyzed the effect of TMEM55B on lysosomal positioning dynamics in isolated primary rat hippocampal neurons. In contrast to non-neuronal cells like ARPE-19 cells (Willett et al., 2017) or HeLa cells (Duhay et al., 2026; Rudnik et al., 2022), we did not observe perinuclear clustering of lysosomes upon the overexpression of TMEM55B or its paralog TMEM55A (**Figure 4A**), suggesting a critical difference between neuronal and non-neuronal cells. The number of axonal lysosomes also did not differ between GFP- and TMEM55B-GFP-transfected cells, as determined by live-cell imaging of LAMP1-RFP expression together with GFP- or TMEM55B-GFP-expressing cells and quantification of the axon tip/axon ratio of LAMP1-positive lysosomes (**Figure 4B**).

**Figure 4.**
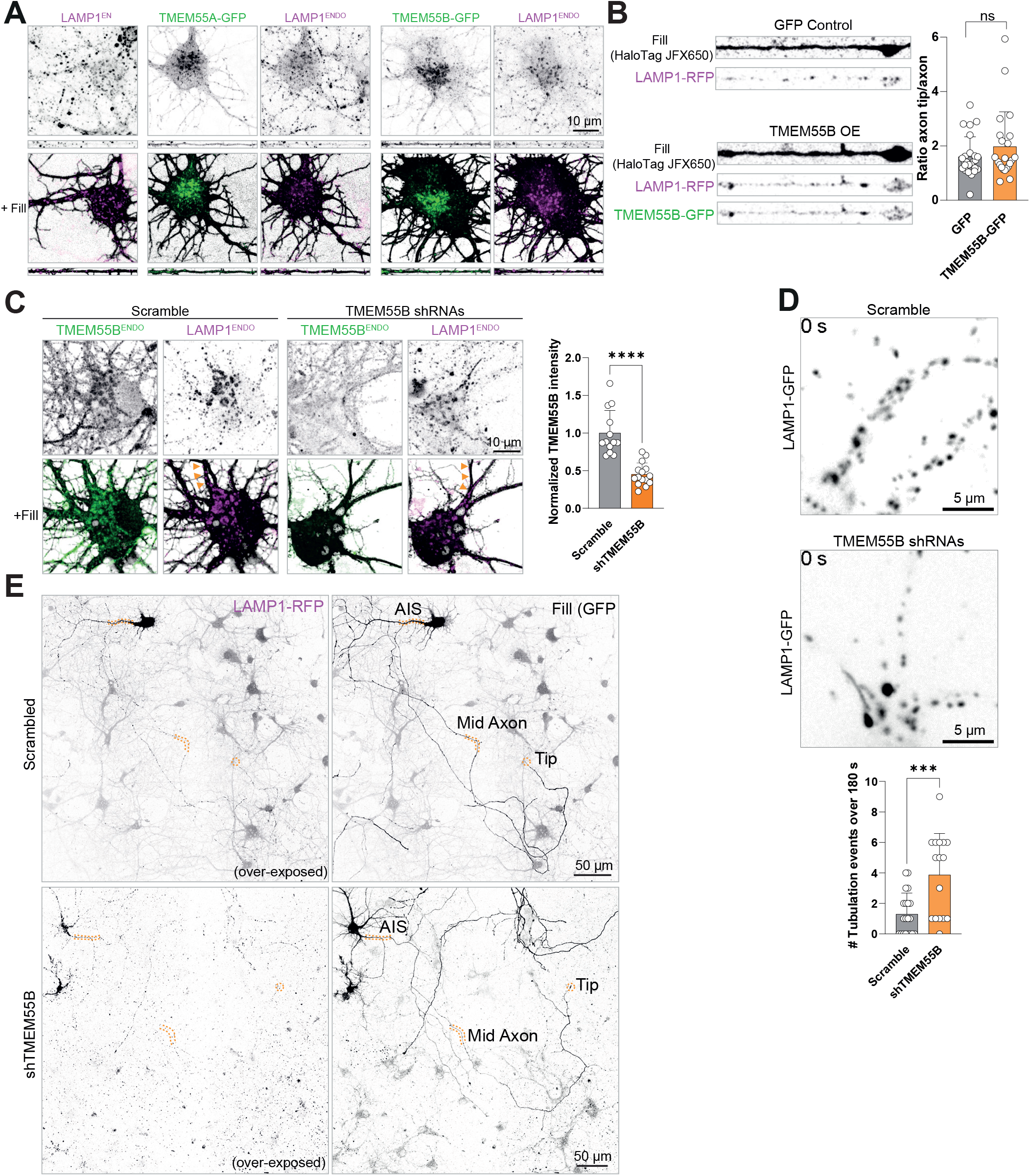
The knockdown of TMEM55B in primary rat neurons leads to lysosomal tubulation defects. **(A)** Immunofluorescence staining for endogenous LAMP1 (LAMP1^endo^, magenta) in rat primary hippocampal neurons overexpressing free GFP, TMEM55A-GFP (green, left), or TMEM55B-GFP (green, right). Cytosolic HaloTag fill (with JFX650 ligand) is shown for visualization of the cell. A representative axon region is shown below. **(B)** Fluorescence microscopy of the axon of rat primary hippocampal neurons expressing LAMP1-RFP together with GFP (upper panel) or TMEM55-GFP (green, lower panel), showing a comparable axonal distribution of LAMP1^+^-lysosomes in the axon. The distribution of LAMP1^+^-lysosomes in the axon was quantified and depicted as the ratio of lysosomes in the axon tip/lysosomes in the axon. Cytosolic HaloTag fill with the JFX650 ligand is shown to visualize the axon. (Unpaired t-test, n=22-25 events; ns = not significant). **(C)** Immunofluorescence staining for endogenous TMEM55B (TMEM55B^endo^) and endogenous LAMP1 in rat primary hippocampal neurons treated with a scramble shRNA or a mixture of two shRNAs targeting TMEM55B. A quantification of the TMEM55B signal intensity is shown (N = 2, n=13-15 cells, unpaired t-test; ns = not significant, **** = p ≤ 0.0001). **(D)** Snapshot of primary rat neurons transfected with LAMP1-GFP and treated with a scramble shRNA or a mixture of shRNAs targeting TMEM55B. Note the tubular structures in the axon of a TMEM55B shRNA-treated neuron. A quantification of the tubular structures observed in live-cell videos over 180 seconds is depicted (N = 3, n=15-20 cells, unpaired t-test; *** = p ≤ 0.001). **(E)** Full view of individual scramble shRNA- or TMEM55B-targeting shRNA neurons transfected with LAMP1-RFP and GFP. (“Fill” / GFP) is shown to outline the transfected cells. The AIS/proximal axon, mid-axon, and axon tip/distal axon are labeled.

To investigate the effect of TMEM55B depletion, we used small hairpin RNA (shRNA). Treating primary rat hippocampal neurons with a combination of two TMEM55B-targeting shRNAs substantially decreased overall TMEM55B signal assessed by immunocytochemistry, indicating effective KD (**Figure 4C**). Interestingly, LAMP1 co-staining revealed tubular LAMP1+-structures at steady-state levels in the axon (**Figure 4C**). This phenotype was more evident in live-cell microscopy experiments, which revealed numerous tubular structures moving dynamically through the proximal axon of TMEM55B shRNA-treated cells (**Figure 4D, Supplemental Video 1**). Quantification of the number of LAMP1^+^-tubular lysosomes in the axon revealed a robust fourfold increase in such structures in shTMEM55B-treated neurons (**Figure 4D**). Overview images of LAMP1-RFP-expressing scramble- and TMEM55B-treated neurons revealed a reduced LAMP signal intensity in the mid- and distal axon (**Figure 4E**).

Next, we investigated the dynamic movement of LAMP1^+^-lysosomes in different axon segments by live-cell imaging in LAMP1-GFP-transfected scramble- and TMEM55B shRNA rat neurons and quantified LAMP1-GFP compartments by dynamic imaging (**Figure 5**). In the proximal axon segment (immediately after the AIS), no differences in the number of anterograde- or retrograde-moving lysosomes or stationary lysosomes were evident (**Figure 5A**, upper panel). In contrast, the number of both anterograde- and retrograde-moving lysosomes was significantly reduced in both the mid-axon and even more clearly in the distal axon (**Figure 5A**, middle and lower panels). The number of stationary lysosomes was similar in the mid-axon but significantly reduced in the distal axon of TMEM55B shRNA-treated neurons. In contrast to the number of dynamic or stationary lysosomes, the velocity of lysosomes did not differ between the mid-axon and the proximal axon (**Figure 5B, Supplemental Figure 6A**). The total number of LAMP1-GFP compartments was unchanged in the proximal axon, significantly reduced in the mid-axon, and even more strongly reduced in the distal axon (**Supplemental Figure 6B**). In conclusion, these data reveal that TMEM55B is critical for the normal distribution of lysosomes in the axon, supporting our *in situ* data in mice. Moreover, our data revealed that both anterograde and retrograde transport are affected by the loss of TMEM55B, consistent with the bidirectional function of JIP4, which binds both kinesins and dynein and mediates bidirectional cargo transport, including lysosomes.

**Figure 5.**
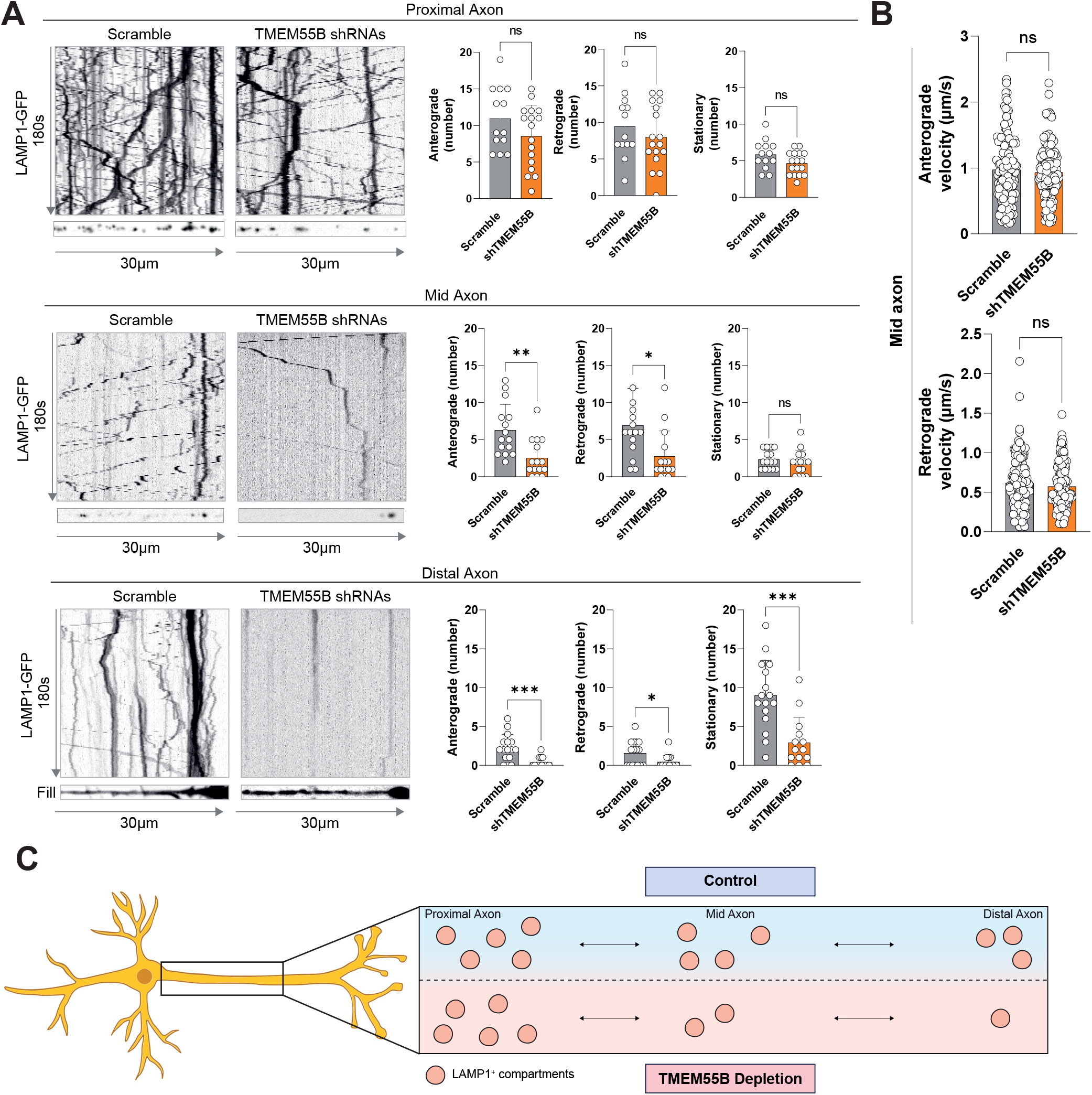
Knockdown of TMEM55B in primary rat neurons affects the bidirectional transport of LAMP1^+^-lysosomes in the mid- and distal axon. **(A)** Representative kymographs and quantifications of LAMP1-GFP-transfected scramble- and TMEM55B shRNA-treated rat primary hippocampal neurons in different axonal subcompartments (upper: proximal axon, middle: mid-axon, and lower: distal axon). The number of anterograde-, retrograde-moving, and stationary lysosomes was quantified. Two-tailed Mann-Whitney test, n=13-16 events; ns = not significant, * = p ≤ 0.05; ** = p ≤ 0.01, *** = p ≤ 0.001). (B) Velocity of LAMP1-GFP-positive organelles moving anterograde (upper graph) and retrograde (lower graph) in the mid-axon. (Two-tailed Mann-Whitney test; n=33-102 events/condition; ns = not significant. (C) Scheme depicting the observed alterations in the distribution of LAMP1+-lysosomes in the different axonal subcompartments due to the depletion of TMEM55B by shRNA.

### Lysosomal dynamics are changed in C. elegans axons depleted of the TMEM55 homolog Y71H2AM.2

To better understand the physiological function of TMEM55B in neurons *in vivo*, we turned to *Caenorhabditis elegans*, which provides a genetically tractable system for *in vivo* imaging. In contrast to mammals and teleost fishes, *C. elegans* and *Drosophila melanogaster* each harbor a single predicted TMEM55 homolog (Y71H2AM.2 and CG6707, respectively) (**Figure 6A**). A multiple sequence alignment of human and mouse TMEM55A and TMEM55B with *C. elegans* Y71H2AM.2 revealed a high degree of homology, similarity, and identity (**Supplemental Figure 7A**). By contrast, the N-terminal region (∼amino acids 1–100) showed the lowest degree of conservation, both between TMEM55A and TMEM55B and relative to Y71H2AM.2. A notable exception was the conserved dileucine-based sorting motif that mediates Golgi-to-lysosome trafficking of TMEM55A and TMEM55B (Rudnik et al., 2022; Willett et al., 2017). The comparatively low conservation of the remaining N-terminal sequence suggests that this region may confer distinct functional properties on TMEM55A and TMEM55B.

**Figure 6.**
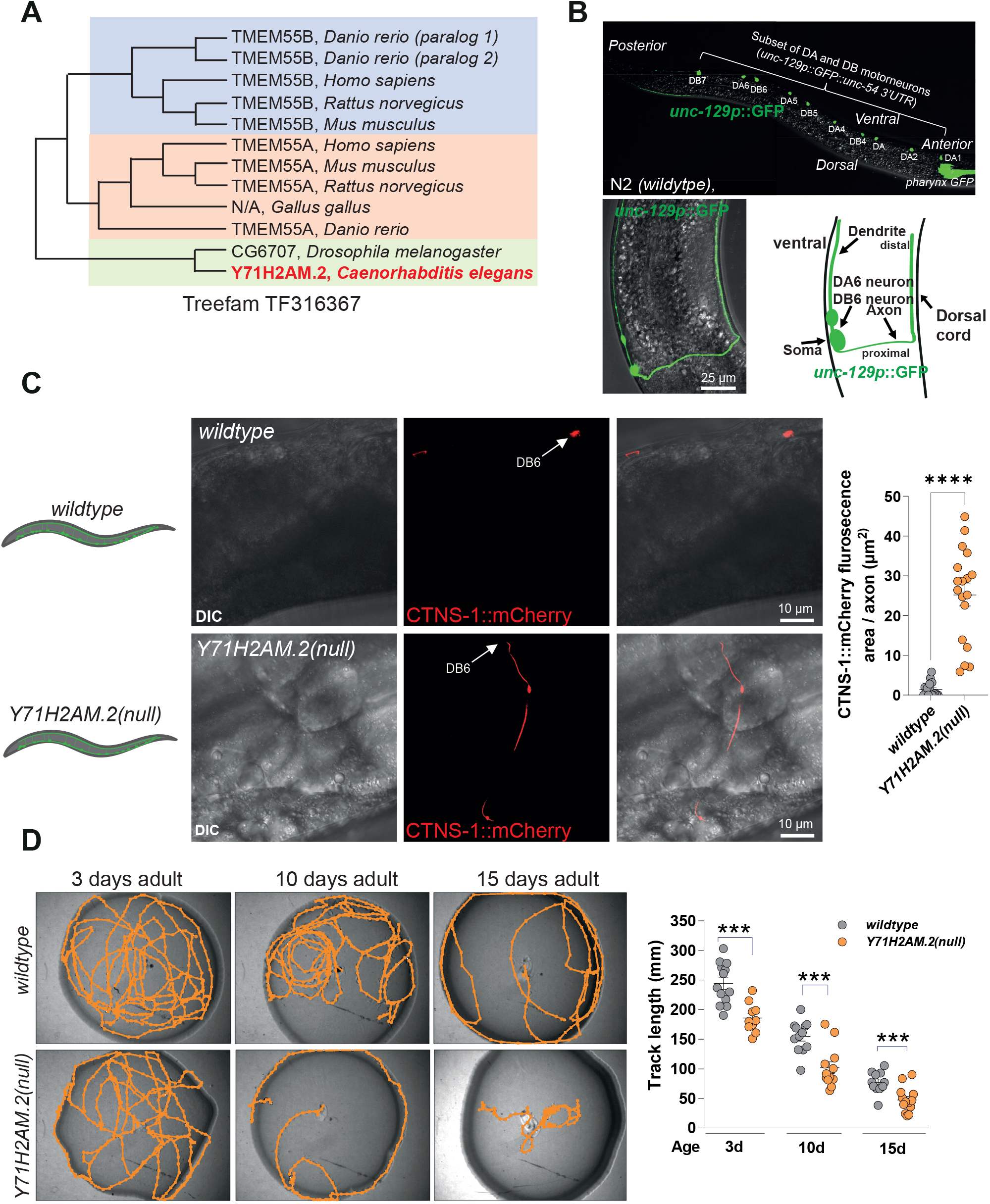
Knockout of the TMEM55B-ortholog Y71H2AM.2 in C. elegans leads to the accumulation of lysosomes in the axon and locomotion deficits. **(A)** Phylogenetic tree inferred from the genomes of Danio rerio, Homo sapiens, Rattus norvegicus, Mus musculus, Drosophila melanogaster, and C. elegans for TMEM55A and TMEM55B and orthologs. (TreeFam, ID: TF316367). (B) Top: Whole-animal fluorescence image showing GFP expression (green) in a subset of cholinergic motor neurons driven by the unc-129(2.64 kb) promoter. Bottom: Live imaging of the DB6 motor neuron expressing unc-129(2.64 kb)p::GFP, showing the DB6 cell body, dendrite, and axon (left panel). A schematic representation of the DB6 motor neuron corresponding to the live image is shown on the right. **(C)** DIC and fluorescence images of N2 wild-type and *Y71H2AM.2(null)* animals expressing the lysosomal marker CTNS-1::mCherry (red) under the control of the *unc-129*(2.64 kb) promoter. Quantification of axonal CTNS-1::mCherry fluorescence in young adult animals (48 h post-hatching) is shown. N = 3 independent experiments, n = 9-10 animals; unpaired t-test; ns, not significant; **** = p ≤ 0.0001. **(D)** Spontaneous locomotor activity of N2 wild-type and *Y71H2AM.2(null)* animals. Crawling distance over 1 h on agarose plates was quantified at 3, 10, and 15 days after synchronization by bleaching. N = 3 independent experiments, n = 10–13 animals per group; one-tailed *t-*test; *** = p ≤ 0.001.

Next, we generated *Y71H2AM.2* KO worms using CRISPR/Cas9-mediated genome editing, hereafter referred to as *Y71H2AM.2(*null) (**Supplemental Figure 7B**). *Y71H2AM.2(null)* worms exhibited a lifespan comparable to that of wildtype animals, indicating that loss of the common TMEM55 ancestor does not affect lifespan under standard conditions (**Supplemental Figure 7C**).

To visualize the cell-specific expression of *Y71H2AM.2*, we first attempted to identify its native promoter sequence within a 4-kb genomic region upstream of the 5’ UTR. However, we could not unambiguously identify it, which indicates that the native *Y71H2AM.2* promoter might be located outside the tested range. As an alternative, we generated worms expressing a cystinosin-mCherry fusion protein (CTNS-1::mCherry) as a lysosomal marker under the control of the *unc-129* promoter (2.64 kb variant) to visualize neural lysosomes *in vivo*. The 2.64-kb *unc-129* promoter fragment drives expression specifically in a subset of DA/DB cholinergic motor neurons (DA1, DA2, DA3, DB4, DA4, DB5, DA5, DB6, DA6, and DB7) located in the *C. elegans* ventral nerve cord (Edwards et al., 2009) (**Figure 6B**). It has been demonstrated that DA and DB motor neurons drive backward and forward locomotion, respectively (Haspel et al., 2010; Wen et al., 2012). We confirmed this expression pattern by fluorescence microscopy using *unc-129*(2.64 kb)p::GFP (**Figure 6B**).

In contrast to wildtype worms, in which lysosomes are excluded from the axon (Niwa et al., 2017) and CTNS-1::mCherry labeling was predominantly confined to motor neuron somata, with weaker signals detected in dendrites, adult *Y71H2AM.2(null)* worms exhibited extensive axonal CTNS-1::mCherry labeling under steady-state conditions (**Figure 6C**). These findings identify Y71H2AM.2 as an important regulator of axonal lysosome positioning.

To determine whether this aberrant axonal lysosome distribution is associated with impaired motor neuron function, we assessed the crawling activity of individual age-matched worms at the ages of 3, 10, and 15 days (**Figure 6D**). At all time points examined, *Y71H2AM.2(null)* worms exhibited a mild but statistically significant reduction in track length compared with N2 wildtype controls. Together, these results indicate that loss of Y71H2AM.2 disrupts the subcellular distribution of CTNS-1::mCherry-positive lysosomes in motor neurons and is accompanied by a mild impairment in locomotor activity.

To establish whether Y71H2AM.2 is a functional homolog of mammalian TMEM55A and TMEM55B, and to examine if the locomotor phenotype of *Y71H2AM.2(null)* animals arises from neuronal rather than muscular dysfunction, we expressed murine TMEM55A or TMEM55B (mTMEM55A and mTMEM55B), each fused to GFP, under the control of the *unc-129*(2.64 kb) promoter. Expression of the lysosomal marker CTNS1::mCherry and mTMEM55B::GFP fusion protein was confirmed by live fluorescence microscopy in the specific subset of motor neurons (**Figures 7A, B, D**). In contrast, expression of mTMEM55A was lethal in both wildtype and *Y71H2AM.2(null)* backgrounds (**Figure 7A**), suggesting that elevated mTMEM55A expression is toxic. While this may reflect supraphysiological expression driven by the *unc-129*(2.64 kb) promoter, we cannot exclude that it indicates a genuine functional divergence between TMEM55A and TMEM55B, which would argue against simple redundancy.

**Figure 7.**
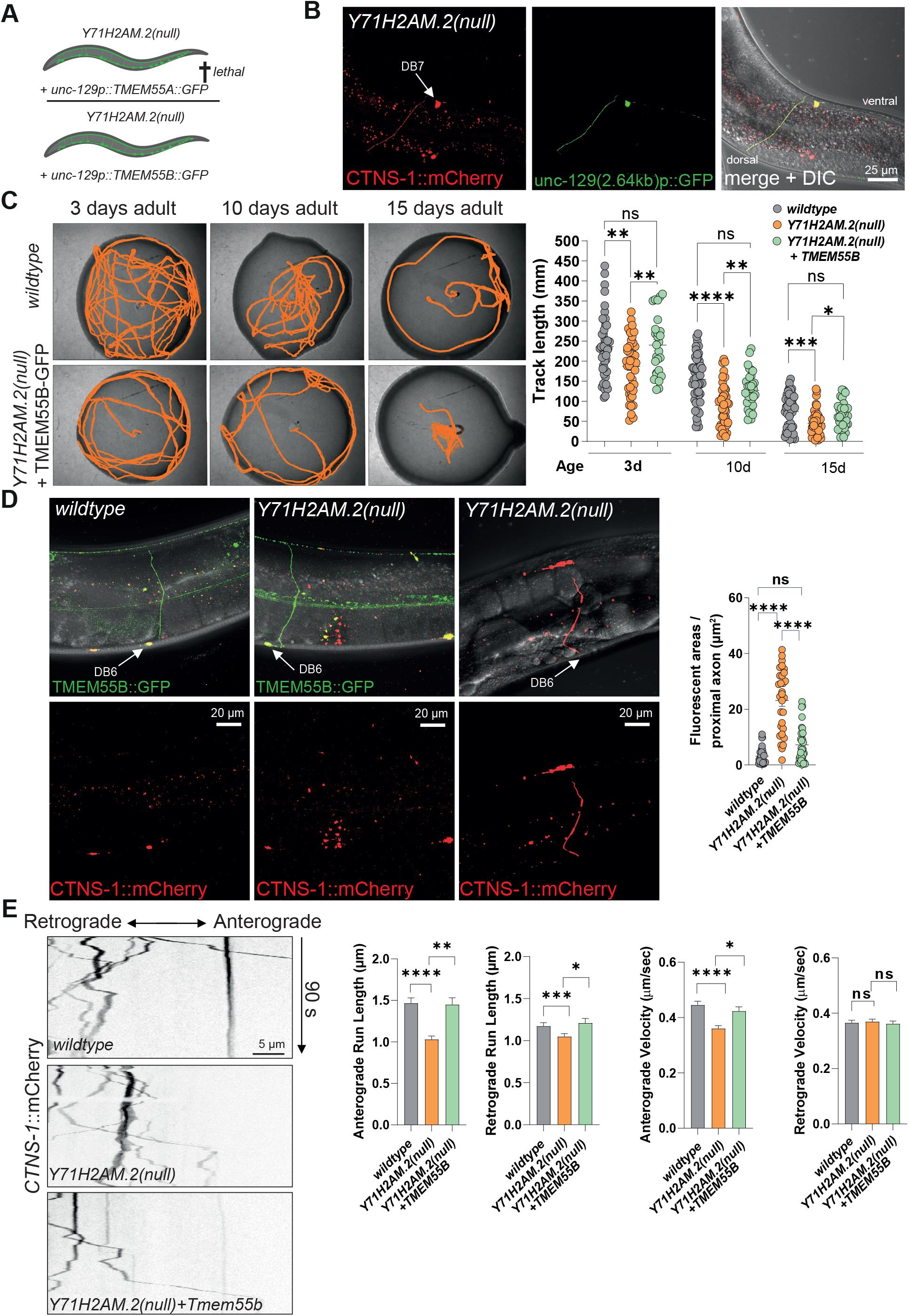
Neuron-specific re-expression of murine TMEM55B in C. elegans Y71H2AM.2(null) animals rescues abnormal lysosomal distribution and locomotor defects. **(A)** Schematic illustrating neuron-specific re-expression of murine TMEM55A-GFP and TMEM55B-GFP in Y71H2AM.2(null) animals. (B) Representative DIC and fluorescence images of DB7 in Y71H2AM.2(null) animals, confirming the specific expression and colocalization of the lysosomal marker CTNS-1::mCherry (red) and GFP (green), both under the control of the unc-129(2.64 kb) promoter. (C) Spontaneous locomotor activity of N2 wild-type, Y71H2AM.2(null), and Y71H2AM.2(null) animals expressing murine TMEM55B (mTMEM55B) in unc-129(2.64 kb)-specific motor neurons. Crawling distance over 1 h on agarose plates seeded with E. coli OP50 was quantified at 3, 10, and 15 days after synchronization by bleaching. N = 3 independent experiments, n = 10–13 animals per group; ordinary one-way ANOVA; * = p ≤ 0.05, ** = p ≤ 0.01, ***= p ≤ 0.001, **** = p ≤ 0.0001. (D) Representative DIC and fluorescence images of young adult (48 h post-hatching) N2 wildtype, Y71H2AM.2(null), and Y71H2AM.2(null)+mTMEM55B-GFP animals expressing unc-129(2.64 kb)p::CTNS-1::mCherry. Axonal CTNS-1::mCherry fluorescence intensity was quantified and compared (right). N ≥ 3 independent experiments, n = 10–11 animals per group; one-way ANOVA with Tukey’s multiple-comparisons test; ns, not significant; **** = p ≤ 0.0001. (E) Representative kymographs of CTNS-1::mCherry-positive lysosome dynamics in the DA9 proximal axon of N2 wild-type, Y71H2AM.2(null), and Y71H2AM.2(null) + mTMEM55B-GFP animals at 3 days after synchronization by bleaching. Anterograde and retrograde run lengths and velocities (individual movement segments) were quantified and compared. n = 731 - 1275 events; Kruskal-Wallis test; ns, not significant; * = p ≤ 0.05, ** = p ≤ 0.01, *** = p ≤ 0.001, **** = p ≤ 0.0001.

By contrast, *Y71H2AM.2(null)* animals expressing mTMEM55B were viable and exhibited a lifespan comparable to that of N2 wildtype animals. Analysis of crawling activity, quantified by track length on agarose plates, revealed that neuronal expression of mTMEM55B fully rescued the locomotor deficit of *Y71H2AM.2(null)* worms at day 3 and resulted in a clear trend toward normalization of locomotor activity at later ages (**Figure 7C**). Moreover, the aberrant accumulation of CTNS-1::mCherry-positive lysosomes in axons of *Y71H2AM.2(null)* worms (**Figure 6C**) was fully reversed by neuronal expression of mTMEM55B under the *unc-129*(2.64 kb) promoter (**Figure 7D**).

In summary, these findings demonstrate that the mammalian TMEM55B can functionally substitute for the *C. elegans* Y71H2AM.2 and support the idea that the locomotor phenotype associated with Y71H2AM.2 loss originates in neurons. This establishes TMEM55B as a functional homolog of Y71H2AM.2 and suggests that TMEM55A and TMEM55B have *in vivo* properties that may differ. Furthermore, rescuing the locomotor phenotype by expressing mTMEM55B specifically in motor neurons under the control of the *unc-129*(2.64 kb) promoter supports a neuronal rather than muscular origin of the motor deficits observed in *Y71H2AM.2(null)* animals.

In order to identify the mechanisms underlying the altered axonal distribution of CTNS-1::mCherry-positive lysosomes and to assess their transport dynamics, we performed *in vivo* imaging in *wildtype*, *Y71H2AM.2(null),* and *Y71H2AM.2(null)* animals expressing mTMEM55B (**Figure 7E**). Notably, analysis of individual movement segments revealed that the loss of Y71H2AM.2 significantly reduced the run length of CTNS-1::mCherry-positive organelles in both anterograde and retrograde directions, with a more pronounced effect on anterograde transport. Motor neuron-specific re-expression of mTMEM55B under the control of the *unc-129*(2.64 kb) promoter fully rescued these defects. In addition, the velocities of CTNS-1::mCherry-positive lysosomes were reduced during anterograde transport but remained unchanged during retrograde transport. Consistent with these findings, analysis of combined trajectories revealed an overall impairment of axonal lysosome transport in *Y71H2AM.2(null)* animals, predominantly affecting the anterograde direction (**Supplemental Figure 7D, E**).

Overall, these results identify Y71H2AM.2 as a regulator of axonal lysosome motility, with a particularly prominent role in anterograde transport, and demonstrate that mammalian TMEM55B can functionally rescue these transport defects *in vivo* in *C. elegans*.

### Tmem55a/b double deficiency leads to impaired growth in mice

To further study TMEM55-dependent lysosomal positioning, we returned to the mouse model and generated *Tmem55a^−/−^* KO mice by CRISPR/Cas9 using a strategy similar to that previously described for *Tmem55b* (**Supplemental Figure 8A-F**). To assess possible functional redundancy, we generated *Tmem55a/b* double KO mice by breeding the two single KO strains. *Tmem55a^−/−^* KO mice were viable and showed no obvious phenotype. *Tmem55a*/b double KO mice, in contrast, were smaller than littermates after weaning. Until they were fully grown, the mice never caught up in weight, with ∼30-50% less weight (**Figure 8A, B**). We observed an increased mortality rate that did not occur at a specific age but rather appeared sporadically (**Figure 8C**). In agreement with the lower body weight, the weight of the brain of *Tmem55a*/b double KO mice was reduced by 25 - 30% compared to wildtype animals (**Figure 8D**). Immunohistochemical survey of the *Tmem55a/b* double KO and a direct comparison with the single KO mice revealed an additional increase in the size of lysosomes, more clumpy aggregation of lysosomes in the area of the PAEZ, and a further increase in the amount of deposited lipofuscin (**Figure 8E**). Microgliosis in the white matter of the spinal cord of *Tmem55a/b^−/−/−-^*dKO mice was comparable with *Tmem55b^−/−^*KO mice and again most pronounced around the ventral medial fissure (**Figure 8F**). Altogether, these results point towards an overall aggravated phenotype in the dKO mice without major effects on neuroinflammation, which is compatible with relatively normal overall development.

**Figure 8.**
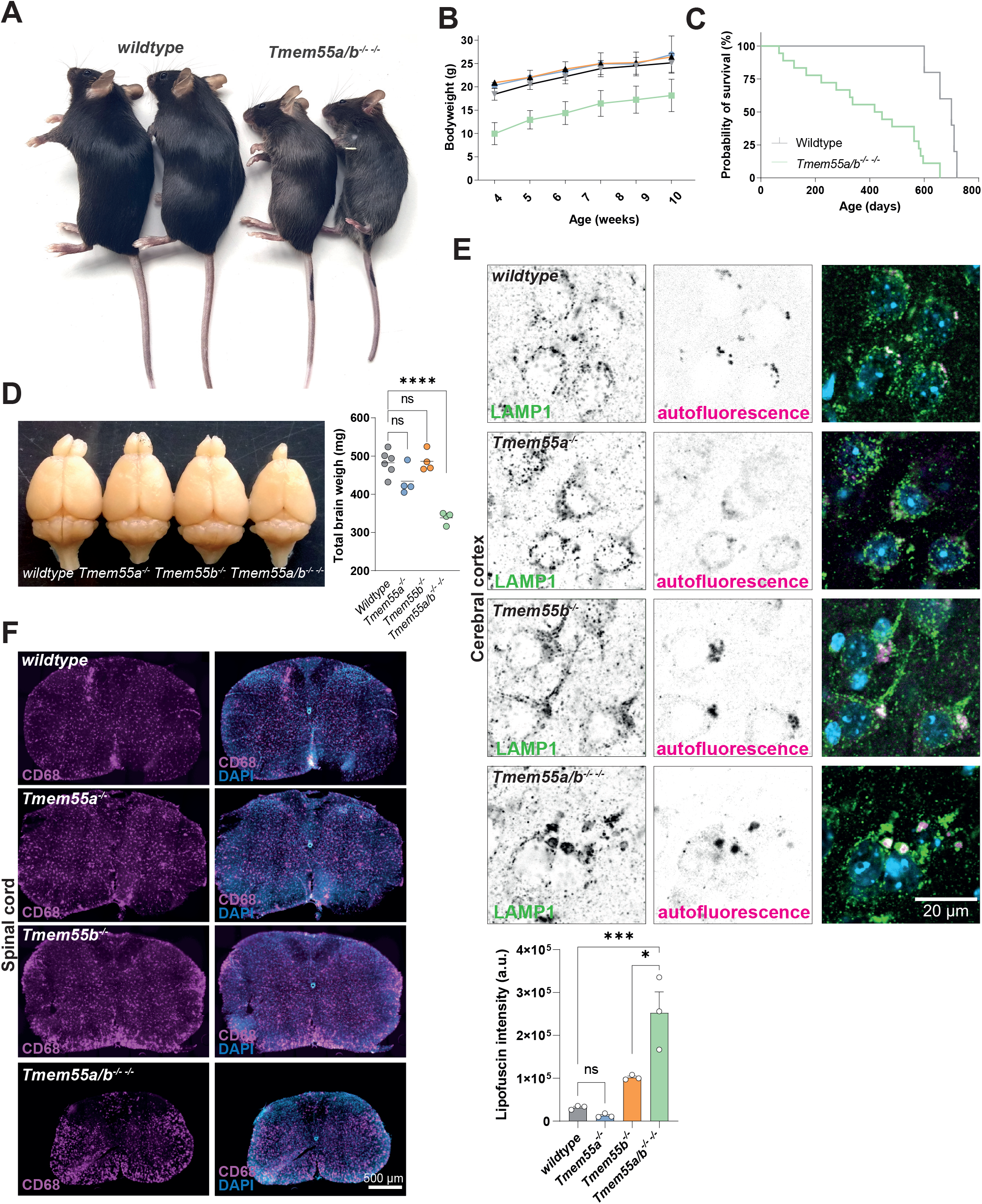
Tmem55a/Tmem55b^−/−−/−^ (Tmem55a/b^−/−−/−^) double KO mice exhibit retarded growth and premature lethality. **(A)** Photograph of two adult (20 weeks) wildtype (left) and two*Tmem55a/b ^−/−−/−^* double KO (dKO) mice (right). **(B)** Bodyweight gain of male wildtype, *Tmem55a^−/−^* KO, *Tmem55b^−/−^* KO, and *Tmem55a/b^−/−−/−^* dKO mice from week 4 – 10 after birth (N = 4-6 animals / genotype). **(C)** Kaplan-Meier survival curve of *wildtype* and *Tmem55a/b^−/−−/−^* mice. **(D)** Photograph of the brains of wildtype, *Tmem55a^−/−^, Tmem55b^−/−^*, and *Tmem55a/b^−/−−/−^*. The total brain weight is depicted (Age: 12 months; N = 3-6 / genotype; One-way ANOVA; ns = not significant, **** = p ≤ 0.0001). **(E)** Immunofluorescence staining of brain sections of wildtype, *Tmem55a^−/^* KO, *Tmem55b^−/−^* KO, and *Tmem55a/b^−/−−/−^* dKO mice with an antibody against LAMP1. The cerebral cortex is shown. Age: 14 months. A quantification of the lipofuscin intensity on sections is shown. N = 3 per genotype, n=50 neurons total; One-way AMOVA; ns = not significant, * = p ≤ 0.05; **** = p ≤ 0.0001). **(F)** Immunofluorescence staining of spinal cord sections from of wildtype, *Tmem55a^−/^* KO, *Tmem55b^−/−^* KO, and *Tmem55a/b^−/−−/−^* dKO mice for CD68 (magenta). Nuclei (blue) are stained with DAPI. Age: 14 months.

## Discussion

Lysosomal positioning is critical in various cell types for the maintenance of cellular homeostasis (Ballabio and Bonifacino, 2020; Bonifacino and Neefjes, 2017). This observation is emphasized by the fact that KO mice for the vast majority of genes that play fundamental roles in this process, in both the retrograde and anterograde directions, are lethal or die immediately after birth. Examples are KOs for the small GTPase Arl8b (Hashimoto et al., 2019), the BORC subunit 7 (BORCS7/Diaskedin, (Snouwaert et al., 2018)), BORC subunit 5 (BORCS5/Myrlysin, (De Pace et al., 2020)), BORC subunit BLOC1S1 (Scott et al., 2014; Zhang et al., 2014), or the retrograde adaptor/motor components Dynein (Harada et al., 1998) or Snapin (Zhou et al., 2011). Interestingly, the *Tmem55b* KO has also been shown to be lethal in mice (Hashimoto et al., 2018). The exact discrepancy between the previous study and ours remains to be clarified. However, based on genomic DNA sequencing validating the CRISPR/Cas9-mediated four-base-pair deletion in the germline and immunoblots, we conclude that our mouse model lacks any remaining TMEM55B protein. Subtle differences in the genetic background, including additional putative modifier genes or differences in methods (CRISPR/Cas9 gene editing vs. gene disruption by a neo/LacZ-containing cassette 57), might explain the apparent differences. Hashimoto et al. identified an interaction of TMEM55B with the mTOR complex 1 (mTORC1) and speculated that the early embryonic lethality is caused by a critical function of TMEM55B in the Rag-Ragulator system on lysosomes. We did not observe any significant differences in amino acid-induced mTORC1 reactivation in MEFs isolated from *Tmem55b^−/−^* KO mice (data not shown), suggesting that TMEM55B is not essential in this process.

The viability of our *Tmem55b^−/−^* KO mouse model allowed us to investigate the physiological function *in vivo* under loss-of-function conditions. A critical finding from our study is the fact that the loss of TMEM55B leads to an apparent lysosomal positioning phenotype even under steady-state physiological conditions in neurons, indicating that the TMEM55B-dependent positioning is more than just a backup mechanism for other adaptor endo-/lysosomal machineries like BORC-, RUFY-, or RAB7/RILP-dependent machineries and that it is not only critical under stress conditions as suggested previously, at least in neurons (Jeong et al., 2024). On the other hand, given the overall mild phenotype, the other above-mentioned mechanisms seem to dominate TMEM55B-dependent lysosomal transport.

Our data revealed a complex interplay between TMEM55B-dependent retro- and anterograde axonal transport of LAMP1^+^-organelles, which is additionally complicated by spatial differences in the axon segment (proximal-, mid-, or distal axon). A key question that still needs extensive additional work is why so many different lysosomes/motor adaptor machineries have developed: Lysosomes are coupled by various proteins and protein complexes to motors (Cabukusta and Neefjes, 2018; Pu et al., 2016), and it is still unclear if they are present on a subpopulation of lysosomes/LAMP1^+^-organelles, used in different cell types, or couple different motors to the same organelles. The change of anterograde and retrograde moving lysosomes in the mid-axon of TMEM55B shRNA-treated neurons is interesting because it could indicate that the few endolysosomes moving in the axon use kinesin-3 instead of kinesin-1, the prototypical motor for lysosomes in the axon (Farias et al., 2017a). These findings were corroborated by our recent study analyzing the axonal transport of LAMP carriers dynamically and support the finding that both kinesin-1 and kinesin-3 are involved in axonal transport, with kinesin-3 being more relevant for fast axonal transport (Li et al., 2024b). There is indeed evidence for sequential motor-driven transport in which kinesin-1 ensures cargo entrance into the AIS and kinesin-3 provides faster transport along the axon (Zahavi et al., 2021). Interestingly, kinesin-3 transports LAMP1 carriers, biosynthetic organelles that deliver lysosomal membrane proteins from the trans-Golgi network to lysosomes (Li et al., 2024a) Notably, they have a typical tubular structure. It is worth speculating that these LAMP1 compartments are not endolysosomes but biosynthetic LAMP carriers. More work is needed to determine the exact motor(s) critical for anterograde, TMEM55B-directed transport, likely mediated via JIP4. The possibility that TMEM55B-organelles could be biosynthetic LAMP carriers could also explain the unexpected finding that only very late and subtle autophagy defects occur in the KO mice, with no changes in LC3-II and only late and relatively little accumulation of p62-positive aggregates, despite the early and primary positioning defects of the LAMP1+-organelles.

We observed *in situ* in *Tmem55b^−/−^* KO mice accumulation of LAMP1^+^-organelles in the PAEZ and the AIS; similarly, we observed tubular structures *in situ* and by live-cell imaging in the very proximal axon of cultured neurons, suggesting a critical role of axon spatial organization. The proximal axon is the critical decision-making site for cargo entry into the axon. Several factors are implicated in this decision, with the AIS having a great effect on the inclusion or exclusion of cargo for axon entry. The PAEZ, on the other hand, was more recently shown to regulate cargo entry to the axon by selectively regulating the attachment of motors to differentially post-translationally modified microtubules (Farias et al., 2015). Our data on the accumulation of LAMP1+ organelles proximal to the axon, compatible with the PAEZ, support a model of impaired loading of lysosomes with anterograde motors such as kinesins. Our data support a hypothesis in which TMEM55B does not control the global abundance of lysosomes, but promotes their transition into a kinesin-competent transport state at the pre-axonal/AIS region. In the absence of TMEM55B, lysosomes accumulate at the axon entry zone, fail to efficiently engage the anterograde transport machinery, and are consequently depleted from distal axonal regions. These findings are in good agreement with the dual role of JIP4, which acts as a scaffold for both dynein and kinesin-1 (Celestino et al., 2022; Vilela et al., 2026). Our findings are also in very good agreement with those from *Jip3/Jip4* double-KO mice. The double-KO mice exhibit premature lethality and swollen axons with an accumulation of typical kinesin-1 cargoes such as APP and synaptophysin (Sato et al., 2015). In addition, a (JIP-dependent) reduction in kinesin-1 binding to microtubules has been demonstrated, with decreased anterograde but not retrograde transport in the axon (Sato et al., 2015). Overall, these findings support the interpretation that anterograde, kinesin-dependent transport is disrupted, rather than dynein-dependent transport.

Although TMEM55B deficiency in mammalian neurons caused an accumulation of LAMP1-positive organelles at the PAEZ/AIS region and depletion from distal axons, loss of the single *C. elegans* homolog resulted in aberrant axonal accumulation of CTNS-1-positive organelles. These findings may reflect species-specific differences in neuronal polarity, the presence and function of the AIS, organelle identity, or the relative contributions of TMEM55-dependent motor-adaptor pathways. It could also could mean that the complexity in its function (in antero- and retrograde direction) was developed later in evolution. Nevertheless, both models support a conserved role in regulating axonal organelle compartmentalization.

The defective binding of JIP4 to lysosomes in *Tmem55b^−/−^* KO mice could also be a decisive factor for the observed preaxonal accumulation of lysosomes. The phenotype of the *Y71H2AM.2(null)* worms interestingly reflects in many regards that of worms depleted of the JIP3/4 homolog UNC-16 (the *C. elegans* homolog of JIP3/4), in particular the redistribution of lysosomes to the more proximal axon (Celestino et al., 2022; Edwards et al., 2015; Edwards et al., 2013), supporting the critical role of the interaction between TMEM55B and JIP4/Y71H2AM.2 and UNC-16. However, UNC-16-deficient worms show a clearly aggravated phenotype, including uncoordinated movement, which is not the case for *Y71H2AM.2(null)* worms, suggesting additional functions of UNC-16 that go beyond its interaction with Y71H2AM.2.

UNC-16 localization, which is concentrated at the initial segment in worms (Edwards et al., 2013), suggesting attachment to cargo in this area. Likewise, JIP3/4 are preferentially attached to LAMP1^+^-autolysosomes in the proximal axon, supporting our hypothesis (Cason and Holzbaur, 2023). The critical dependence of TMEM55B on JIP4 is further substantiated by the phenocopy of the *Jip4* cKO and *Tmem55b* KO in neurons on lysosomal positioning. It is interesting that JIP4 deficiency (likely in complex with TMEM55B) alone has such a strong effect in neurons, given that JIP3 seems to be even more critical for axonal transport, and they have been shown to have at least partially redundant functions (Gowrishankar et al., 2021). These data also highlight the complexity among different adaptor machineries, providing evidence of partial, but not fully overlapping, functions.

To investigate potential redundancy between TMEM55A and TMEM55B, we generated double KO mice. We did not observe any phenotype in *Tmem55a^−/−^*KO mice. The *Tmem55a/b^−/−−/−^* dKO mice showed reduced weight gain; surprisingly, they did not make up the weight as they aged. In addition, the animals showed a significantly increased mortality rate, with ∼50% dying within the first year. We could not determine the cause of their death, and, surprisingly, the neuronal lysosomal positioning phenotype was slightly exacerbated. Neurons showed larger, more densely clustered lysosomes in the PAEZ, but neuroinflammation was not exacerbated in the spinal cord. These findings do not distinguish between functional redundancy and additive, independent effects, as they leave open the possibility that both proteins have a redundant function and are both critical in a particular organ, for example. We may also have overlooked a phenotype in *Tmem55a^−/−^* KO mice, and the additive effects of TMEM55A and TMEM55B deficiency may explain the growth and survival phenotypes. However, given the direct effect on lysosomal clustering, our data suggest a previously uncharacterized functional redundancy and a role for TMEM55A in lysosomal positioning.

The crystal structure of the cytosolic region covering residues 80-166 of TMEM55B has recently been solved alone and in complex with a C-terminal RILPL1 peptide (Waschbusch et al., 2026), underscoring its relevance in retrograde transport. However, additional interaction partners were identified, among them JIP4 and TBC1D9B, which we found independently as a TMEM55B-binder that acts as a GTPase-activating protein (GAP) for ARL8B (Duhay et al., 2026), which is also critical for anterograde lysosomal positioning, e.g., by regulating the binding of BORC to lysosomes. More work is needed to distinguish TMEM55B’s role in mediating these functions. In particular, the interplay between JIP4 Binding and TBC1D9B may be mutually exclusive or competitive and warrants future study.

In summary, our work extends the knowledge of TMEM55B on lysosomal transport *in vivo* in highly polarized neuronal cells and provides robust evidence that TMEM55B has a bifunctional role in both retrograde and notably anterograde transport of lysosomes, which likely depends mostly on JIP4 / UNC-16 in *C. elegans*. Moreover, surprisingly, TMEM55A and TMEM55B seem to play at least partially redundant functions in the mouse, although additive effects can not be entirely excluded.

## Acknowledgments

We thank Sebastian Held, Dana Germer, Annika Detje, and Jana Kopkanova for their excellent technical assistance. Julia Bär and Marina Mikhaylova are acknowledged for initial help with primary neurons. The authors declare no competing financial or non-financial interests. The Czech Centre for Phenogenomics was supported by the Czech Academy of Sciences (RVO 68378050) and by the Ministry of Education, Youth and Sports of the Czech Republic through project LM202303, Czech Centre for Phenogenomics, to R.S. This work was supported by the German Research Foundation (DFG) with grant SA 683/9-1 (FOR2625) to P.S. and DA 1785/2-1 and DA 1785/2-2 to M.D. (FOR2625).

## Data availability

The transcriptomic data generated from NanoString nCounter panels and RNA sequencing in this study are provided as Supplementary Information files. Uncropped blots are also presented in the Supplementary Information. All other data supporting the findings of this study are included within the main text of the article. Source data are provided with this paper.

## Author contributions

S.R., C.H.L., D.-C.G., A.V., D.J.B., J.R., S.St., B.S., I.M.B., R.H. and M.R. performed experiments and acquired data. K.Y. provided essential material (JIP4 conditional knockout brains). R.S., F.D., P.S., R.D., R.G., G.G.F. and M.D. contributed to data analysis and interpretation and/or supervised the work. M.D. conceived the study and wrote the manuscript; all authors critically read and approved the final version.

## Abbreviations

DAPI: 4’,6-Diamidino-2-phenylindole
AIS: axon initial segment
BORC: BLOC-one-related complex
CRISPR: Clustered Regularly Interspaced Short Palindromic Repeats
PI: phosphatidylinositol
LE: late endosomes
OPCs: oligodendrocyte progenitor cells
PAEZ: pre-axonal exclusion zone
KD: knockdown
KO: knockout
shRNA: small hairpin RNA

## SUPPLEMENTAL FIGURE LEGENDS

**Supplemental Figure 1. *TMEM55B is highly expressed in neurons of the mouse brain.* (A)** Immunoblot of multiple tissue extracts from an adult mouse (6 months) with an antibody against TMEM55B. Actin and GAPDH are depicted as loading controls. Quantification of the TMEM55B signal across different tissues from N = 3 animals is shown. **(B)** Overview image of the cerebral cortex from an immunofluorescence staining of the mouse brain with an antibody against TMEM55B (white) and DAPI-stained nuclei (blue). (Age: 3 months). **(C)** Immunofluorescence staining of the mouse brain with antibodies against TMEM55B (red) and cell-specific markers (green), NeuN (neurons), GFAP (astrocytes), Iba1 (microglia), and Ng2 (oligodendrocyte progenitor cells); Nuclei are stained with DAPI (blue). **(D)** Immunofluorescence staining of the mouse brain with antibodies against LAMP1 (green) and TMEM55B (red); Nuclei are stained with DAPI (blue). A cortical neuron is depicted. C, D: Age: 3 months.

**Supplemental Figure 2. *oGeneration f Tmem55b^−/−^ KO mice by CRISPR/Cas9.* (A)** Exon/intron structure of the *Tmem55b* gene in mice. The site of the CRISPR/Cas9-edited exon (exon 2) is indicated. **(B)** Sequence of the *Tmem55b*-targeting CRISPR guide in exon two (grey box). The PAM sequence (CCC) is highlighted in orange. The coded amino acid sequence is depicted below. **(C)** Mutations obtained from gene editing as assessed by Sanger sequencing (Del = deletion). The mouse line chosen for further breeding and experiments in this manuscript is boxed in red. **(D)** Sanger sequencing chromatogram of exon 2 of the chosen mouse line containing a 4-nucleotide (nt) deletion. **(E)** High-resolution capillary electrophoresis chromatogram of the genotyping PCR product used for genotyping of a *Tmem55b^+/+^*, a heterozygous *Tmem55b^+/−^*, and a homozygous *Tmem55b^−/−^* mouse. **(F)** Genotype distribution of the offspring of heterozygous *Tmem55b^+/−^* x *Tmem55b^+/−^*breeding couples. **(G)** Immunoblot of tissue lysates and mouse embryonic fibroblasts (MEFs) from the chosen *Tmem55b^−/−^* KO mouse line (4 nt deletion, (#36607)) with an antibody against TMEM55B. GAPDH is shown as a loading control.

**Supplemental Figure 3. *Lysosomal markers and the autophagy marker LC3-II are unchanged in Tmem55b^−/−^ KO mice.* (A)** Principal component analysis (PCA) of bulk RNA sequencing data from total brain RNA of wildtype (WT1 - 4) and *Tmem55b*^−/−^ KO (KO1 - 4) mice, showing a clear separation of the two genotype groups along principal component 1. Age: 10 months. **(B)** Immunoblot analysis of total brain lysates from *Tmem55B^+/+^* and *Tmem55b^−/−^* KO mice for Cathepsin D (CTSD), LC3, LAMP1, TMEM55B, and GAPDH (loading control). The apparent molecular weights of LC3-I and LC3-II are indicated by arrowheads. Age: 16 months.

**Supplemental Figure 4. *Behavioral survey of Tmem55b^−/−^ mice.* (A)** *Tmem55b^+/+^* and *Tmem55b^−/−^* KO mice showed no difference in grip strength (t = 1.1, p = 0.28). **(B)** Mice generally showed improved performance over sequential rotarod trials (F = 9.3, p < 0.001). *Tmem55b^−/−^* KO mice tended to perform better than wildtype controls (F = 3.3, p = 0.08), independent of trials (F = 0.2, p = 0.92). An ANOVA was performed comparing genotypes for average drop latency, with weight as a covariate. Weight significantly influenced drop latency (p < 0.01), but there was no significant difference between the genotypes (p = 0.22). **(C)** Mice were ventrally video-tracked during treadmill walking to allow parameter extraction for gait analysis. Fore vs hind gait symmetry (the ratio of forelimb stepping frequency to hind limb stepping frequency) was similar in both genotypes (U = 85.5, p = 0.58). Likewise, there were no significant differences in spatial parameters including fore and hind stance widths (gen F = 3.2, p = .08 & gen x paws F = 1.8, p = 0.19), step angles (gen F = 0.0, p = 0.92 & gen x paws F = 0.2, p = 0.64) or stride lengths (gen F = 0.0, p = 0.85 & gen x paws F = 0.5, p = 0.51). The average stride time (including stance and swing times) was not different between the genotypes (gen F = 0.0, p = 0.87 & gen x paws F = 0.5, p = 0.50). **(D)** *Tmem55B^+/+^* and *Tmem55b^−/−^* KO mice showed no difference in open field behavior. Both genotypes traveled similar distances during 10 min of free exploration (t = 0.8, p = 0.44). Time spent in the center (t = 0.5, p = 0.61) and time spent in the periphery (t = 0.5, p = 0.65) were comparable as well. Heat maps illustrate a similar distribution of positions for both genotypes. **(E)** *Tmem55B^+/+^* and *Tmem55b^−/−^*KO mice showed no differences in latency to traverse the square or round beams (F = 0.2, p = 0.70; F = 0.1, p = 0.77). Likewise, no differences were observed in the number of paw slips on square or round beams (F = 2.5, p = 0.13; F = 1.4, p = 0.25). Genotypes were compared for average latency and number of slips using weight as a covariate. Weight did not significantly affect latency, and adjusted means were not different between genotypes (p = 0.67). **(F)** During 23 h cage activity recording, mice showed characteristic nocturnal activity peaks (effect of time point: F = 24.3, p < 0.001, (2-way RM ANOVA)). *Tmem55b^−/−^* KO appeared to be more active than wildtype controls (F = 5.8, p < 0.05), but this effect depended on the time point (F = 1.6, p < 0.01). Specifically, *Tmem55b^−/−^*KO mice were more active during the first nocturnal peak (between 20 h and 23.30 h). *Tmem55b^−/−^* KO mice remained more active during this nocturnal peak after taking into account weight as a covariate (p < 0.05). **(G)** There was no difference between *Tmem55B^+/+^*and *Tmem55b^−/−^* KO mice for latency to tail withdrawal (U = 88, p = 0.62), suggesting similar responsivity to a nociceptive stimulus. **(H)** *Tmem55B^+/+^* and *Tmem55b^−/−^* KO mice showed similar total activity levels in the elevated plus maze (t = 1.3, p = 0.19). There was no significant difference in the percentage of beam crossings in the open arms (t = 1.2, p = 0.25). *Tmem55b^−/−^* KO mice tended to spend relatively less time in the open arms (t = 2.0, p = 0.06).

**Supplemental Figure 5. *Tissue expression of JIP4 in mice and normal distribution of lysosomes in forebrain conditional Jip4 KO mice in the thalamus.* (A)** Immunoblot analysis of adult mouse tissues with a JIP4-specific antibody (age: 10 months). Actin is depicted as a loading control. Densitometric quantification of immunoblots of N = 3 animals of different tissues for JIP4 signals. (B) Immunofluorescence staining of brain sections of floxed EMX cre-negative JIP4 mice (*JIP4^fl/fl^* EMX cre-) and conditional forebrain-specific *Jip4* KO mice (*JIP4^fl/fl^* EMX cre+) for LAMP1 (green) and autofluorescence (red) of the thalamus. Note the normal distribution of lysosomes in the thalamus, where the EMX promoter is not active. Age: 3 months.

**Supplemental Figure 6. *Velocity of LAMP1-GFP organelles in the proximal axon of TMEM55B shRNA-treated primary rat neurons.* (A)** Velocity of LAMP1-GFP-positive organelles moving anterograde (upper graph) and retrograde (lower graph) in the proximal axon. (Two-tailed Mann-Whitney test; n = 33-102 events/condition; ns = not significant. **(B)** Quantification of the total number of LAMP1-GFP-positive organelles in the proximal-, mid-, and distal axon. (Unpaired t-test; N = 3; n = 13 - 17 cells; ns = not significant., ** = p ≤ 0.01, **** = p ≤ 0.0001).

**Supplemental Figure 7. *Alignment of TMEM55A and TMEM55B and homologs from different species and generation of Y71H2AM.2 KO worms by CRISPR/Cas9.* (A)** Sequence logo of Y71H2AM.2, human (h)TMEM55B, mouse (m)TMEM55B, hTMEM55A, and mTMEM55A. (B) Agarose gel of genotyping PCRs for *Y71H2AM.2(null)* background in the wildtype N2 and transgenes. The PCR primers amplify *Y71H2AM.2* in the N2 wildtype (+/+; 1020 bp), the *Y71H2AM.2(null)* mutants (−/−), and transgenic null mutants carrying the extrachromosomal constructs *CTNS-1::mCherry*, *TMEM55B::GFP*, or both in the *Y71H2AM.2(null)* background (−/−; 1290 bp). (C) Survival curve of wildtype and *Y71H2AM.2(null)* worms kept at 20 °C. (D) Quantification of anterograde- and retrograde run length of CTNS-1::mCherry-positive lysosomes in the DA9 axon of wildtype, *Y71H2AM.2(null)*, and *Y71H2AM.2(null)* + mTMEM55B-GFP worms (Age: 3 days (larvae)). The combined trajectories were quantified (n=507-824 events, Kruskal-Wallis test, ns = not significant, ** = p ≤ 0.01, *** = p ≤ 0.001). (E) Quantification of anterograde- and retrograde velocity of *ctns-1::*mCherry-positive lysosomes in the DA9 axon of wildtype, *Y71H2AM.2(null),* and *Y71H2AM.2(null)* + mTMEM55B-GFP worms (Age: 3 days (larvae)). Combined trajectories were quantified (n = 507-824 events; Kruskal-Wallis test; ns = not significant; **** = p ≤ 0.0001).

**Supplemental Figure 8. *Generation of Tmem55a^−/−^ KO mice by CRISPR/Cas9.* (A)** Exon/intron structure of the *Tmem55a* gene in mice. The site of the CRISPR/Cas9-edited exon (exon 2) is indicated. **(B)** Sequence of the *Tmem55a*-targeting CRISPR guide in exon two (grey box). The PAM sequence (CCC) is highlighted in red. The coded amino acid sequence is depicted below. **(C)** Mutations obtained from gene editing (Del = deletion) as assessed by Sanger sequencing. The mouse line chosen for further breeding and experiments is boxed in red. **(D)** Sanger sequencing chromatogram of exon two of the chosen line containing an 8-nucleotide (nt) deletion. **(E)** High-resolution capillary electrophoresis chromatogram of the genotyping PCR product used for genotyping of a *Tmem55a^+/+^*, a heterozygous *Tmem55a^+/−^*, and a homozygous *Tmem55a^−/−^* KO mouse. **(F)** Genotype distribution of the offspring of heterozygous *Tmem55b*^+/−^ x *Tmem55a^+/−^*breeding couples.

## References

1. Adnan, G., Rubikaite, A., Khan, M., Reber, M., Suetterlin, P., Hindges, R., and Drescher, U. (2020). The GTPase Arl8B Plays a Principle Role in the Positioning of Interstitial Axon Branches by Spatially Controlling Autophagosome and Lysosome Location. J Neurosci 40, 8103–8118.

2. Anders, S., Pyl, P.T., and Huber, W. (2015). HTSeq--a Python framework to work with high-throughput sequencing data. Bioinformatics 31, 166–169.

3. Ballabio, A., and Bonifacino, J.S. (2020). Lysosomes as dynamic regulators of cell and organismal homeostasis. Nat Rev Mol Cell Biol 21, 101–118.

4. Becker, N.H., Goldfischer, S., Shin, W.Y., and Novikoff, A.B. (1960). The localization of enzyme activities in the rat brain. J Biophys Biochem Cytol 8, 649–663.

5. Bolstad, B.M., Irizarry, R.A., Astrand, M., and Speed, T.P. (2003). A comparison of normalization methods for high density oligonucleotide array data based on variance and bias. Bioinformatics 19, 185–193.

6. Bonifacino, J.S., and Neefjes, J. (2017). Moving and positioning the endolysosomal system. Curr Opin Cell Biol 47, 1–8.

7. Brenner, S. (1974). The genetics of Caenorhabditis elegans. Genetics 77, 71–94.

8. Cabukusta, B., and Neefjes, J. (2018). Mechanisms of lysosomal positioning and movement. Traffic 19, 761–769.

9. Cason, S.E., and Holzbaur, E.L.F. (2022). Selective motor activation in organelle transport along axons. Nat Rev Mol Cell Biol.

10. Cason, S.E., and Holzbaur, E.L.F. (2023). Axonal transport of autophagosomes is regulated by dynein activators JIP3/JIP4 and ARF/RAB GTPases. J Cell Biol 222.

11. Cason, S.E., Mogre, S.S., Holzbaur, E.L.F., and Koslover, E.F. (2022). Spatiotemporal analysis of axonal autophagosome-lysosome dynamics reveals limited fusion events and slow maturation. Mol Biol Cell, mbcE22030111.

12. Cataldo, A.M., and Nixon, R.A. (1990). Enzymatically active lysosomal proteases are associated with amyloid deposits in Alzheimer brain. Proc Natl Acad Sci U S A 87, 3861–3865.

13. Celestino, R., Gama, J.B., Castro-Rodrigues, A.F., Barbosa, D.J., Rocha, H., d’Amico, E.A., Musacchio, A., Carvalho, A.X., Morais-Cabral, J.H., and Gassmann, R. (2022). JIP3 interacts with dynein and kinesin-1 to regulate bidirectional organelle transport. J Cell Biol 221.

14. Cheng, X.T., Xie, Y.X., Zhou, B., Huang, N., Farfel-Becker, T., and Sheng, Z.H. (2018). Characterization of LAMP1-labeled nondegradative lysosomal and endocytic compartments in neurons. J Cell Biol 217, 3127–3139.

15. Damme, M., and Kummel, D. (2026). TMEM55B-Jack of all trades in the endo-lysosomal system? Structure 34, 215–217.

16. Damme, M., Stroobants, S., Walkley, S.U., Lullmann-Rauch, R., D’Hooge, R., Fogh, J., Saftig, P., Lubke, T., and Blanz, J. (2011). Cerebellar alterations and gait defects as therapeutic outcome measures for enzyme replacement therapy in alpha-mannosidosis. J Neuropathol Exp Neurol 70, 83–94.

17. De Pace, R., Britt, D.J., Mercurio, J., Foster, A.M., Djavaherian, L., Hoffmann, V., Abebe, D., and Bonifacino, J.S. (2020). Synaptic Vesicle Precursors and Lysosomes Are Transported by Different Mechanisms in the Axon of Mammalian Neurons. Cell Rep 31, 107775.

18. Duhay, V., Tian, M., Kosieradzka, K., Ebner, M., Lo, W.T., Krauss, M., Sprengel, H.L., Voss, M., Riechmann, M., Savas, J.N., et al. (2026). Control of lysosome function by the GTPase-activating protein TBC1D9B and its binding partner TMEM55B. Nat Commun 17.

19. Edwards, S.L., Charlie, N.K., Richmond, J.E., Hegermann, J., Eimer, S., and Miller, K.G. (2009). Impaired dense core vesicle maturation in Caenorhabditis elegans mutants lacking Rab2. J Cell Biol 186, 881–895.

20. Edwards, S.L., Morrison, L.M., Yorks, R.M., Hoover, C.M., Boominathan, S., and Miller, K.G. (2015). UNC-16 (JIP3) Acts Through Synapse-Assembly Proteins to Inhibit the Active Transport of Cell Soma Organelles to Caenorhabditis elegans Motor Neuron Axons. Genetics 201, 117–141.

21. Edwards, S.L., Yu, S.C., Hoover, C.M., Phillips, B.C., Richmond, J.E., and Miller, K.G. (2013). An organelle gatekeeper function for Caenorhabditis elegans UNC-16 (JIP3) at the axon initial segment. Genetics 194, 143–161.

22. Farfel-Becker, T., Roney, J.C., Cheng, X.T., Li, S., Cuddy, S.R., and Sheng, Z.H. (2019). Neuronal Soma-Derived Degradative Lysosomes Are Continuously Delivered to Distal Axons to Maintain Local Degradation Capacity. Cell Rep 28, 51–64 e54.

23. Farias, G.G., Guardia, C.M., Britt, D.J., Guo, X., and Bonifacino, J.S. (2015). Sorting of Dendritic and Axonal Vesicles at the Pre-axonal Exclusion Zone. Cell Rep 13, 1221–1232.

24. Farias, G.G., Guardia, C.M., De Pace, R., Britt, D.J., and Bonifacino, J.S. (2017a). BORC/kinesin-1 ensemble drives polarized transport of lysosomes into the axon. Proc Natl Acad Sci U S A 114, E2955–E2964.

25. Farias, G.G., Guardia, C.M., De Pace, R., Britt, D.J., and Bonifacino, J.S. (2017b). BORC/kinesin-1 ensemble drives polarized transport of lysosomes into the axon. Proc Natl Acad Sci U S A 114, E2955–E2964.

26. Gottschling, D.C., Doring, F., and Luersen, K. (2017). Locomotion Behavior Is Affected by the Galpha(S) Pathway and the Two-Pore-Domain K(+) Channel TWK-7 Interacting in GABAergic Motor Neurons in Caenorhabditis elegans. Genetics 206, 283–297.

27. Gowrishankar, S., Lyons, L., Rafiq, N.M., Roczniak-Ferguson, A., De Camilli, P., and Ferguson, S.M. (2021). Overlapping roles of JIP3 and JIP4 in promoting axonal transport of lysosomes in human iPSC-derived neurons. Mol Biol Cell 32, 1094–1103.

28. Guardia, C.M., Farias, G.G., Jia, R., Pu, J., and Bonifacino, J.S. (2016). BORC Functions Upstream of Kinesins 1 and 3 to Coordinate Regional Movement of Lysosomes along Different Microtubule Tracks. Cell Rep 17, 1950–1961.

29. Harada, A., Takei, Y., Kanai, Y., Tanaka, Y., Nonaka, S., and Hirokawa, N. (1998). Golgi vesiculation and lysosome dispersion in cells lacking cytoplasmic dynein. J Cell Biol 141, 51–59.

30. Hashimoto, K., Yamaguchi, Y., Kishi, Y., Kikko, Y., Takasaki, K., Maeda, Y., Matsumoto, Y., Oka, M., Miura, M., Ohata, S., et al. (2019). Loss of the small GTPase Arl8b results in abnormal development of the roof plate in mouse embryos. Genes Cells 24, 436–448.

31. Hashimoto, Y., Shirane, M., and Nakayama, K.I. (2018). TMEM55B contributes to lysosomal homeostasis and amino acid-induced mTORC1 activation. Genes Cells 23, 418–434.

32. Haspel, G., O’Donovan, M.J., and Hart, A.C. (2010). Motoneurons dedicated to either forward or backward locomotion in the nematode Caenorhabditis elegans. J Neurosci 30, 11151–11156.

33. Jeong, E., Willett, R., Rissone, A., La Spina, M., and Puertollano, R. (2024). TMEM55B links autophagy flux, lysosomal repair, and TFE3 activation in response to oxidative stress. Nat Commun 15, 93.

34. Jungnickel, K.E.J., Guelle, O., Iguchi, M., Dong, W., Kotov, V., Gabriel, F., Debacker, C., Dairou, J., McCort-Tranchepain, I., Laqtom, N.N., et al. (2024). MFSD1 with its accessory subunit GLMP functions as a general dipeptide uniporter in lysosomes. Nat Cell Biol 26, 1047–1061.

35. Kechin, A., Boyarskikh, U., Kel, A., and Filipenko, M. (2017). cutPrimers: A New Tool for Accurate Cutting of Primers from Reads of Targeted Next Generation Sequencing. J Comput Biol 24, 1138–1143.

36. Kenyon, C. (1988). The nematode Caenorhabditis elegans. Science 240, 1448–1453.

37. Kenyon, C., Chang, J., Gensch, E., Rudner, A., and Tabtiang, R. (1993). A C. elegans mutant that lives twice as long as wild type. Nature 366, 461–464.

38. Li, C.H., Kersten, N., Ozkan, N., Nguyen, D.T.M., Koppers, M., Post, H., Altelaar, M., and Farias, G.G. (2024a). Spatiotemporal proteomics reveals the biosynthetic lysosomal membrane protein interactome in neurons. Nat Commun 15, 10829.

39. Li, C.H., Kersten, N., Ozkan, N., Nguyen, D.T.M., Koppers, M., Post, H., Altelaar, M., and Farias, G.G. (2024b). Spatiotemporal proteomics reveals the biosynthetic lysosomal membrane protein interactome in neurons. Nat Commun 15, 10829.

40. Lie, P.P.Y., Yang, D.S., Stavrides, P., Goulbourne, C.N., Zheng, P., Mohan, P.S., Cataldo, A.M., and Nixon, R.A. (2021). Post-Golgi carriers, not lysosomes, confer lysosomal properties to pre-degradative organelles in normal and dystrophic axons. Cell Rep 35, 109034.

41. Liu, X., Dos Santos, T., Spigelman, A.F., Duckett, S., Smith, N., Suzuki, K., and MacDonald, P.E. (2025). TMEM55A-mediated PI5P signalling regulates alpha cell actin depolymerisation and glucagon secretion. Diabetologia.

42. Luersen, K., Gottschling, D.C., and Doring, F. (2016). Complex Locomotion Behavior Changes Are Induced in Caenorhabditis elegans by the Lack of the Regulatory Leak K+ Channel TWK-7. Genetics 204, 683–701.

43. Maday, S., Wallace, K.E., and Holzbaur, E.L. (2012). Autophagosomes initiate distally and mature during transport toward the cell soma in primary neurons. J Cell Biol 196, 407–417.

44. Montagnac, G., Sibarita, J.B., Loubery, S., Daviet, L., Romao, M., Raposo, G., and Chavrier, P. (2009). ARF6 Interacts with JIP4 to control a motor switch mechanism regulating endosome traffic in cytokinesis. Curr Biol 19, 184–195.

45. Morioka, S., Nigorikawa, K., Okada, E., Tanaka, Y., Kasuu, Y., Yamada, M., Kofuji, S., Takasuga, S., Nakanishi, H., Sasaki, T., et al. (2018). TMEM55a localizes to macrophage phagosomes to downregulate phagocytosis. J Cell Sci 131.

46. Nguyen, Q., Lee, C.M., Le, A., and Reddy, E.P. (2005). JLP associates with kinesin light chain 1 through a novel leucine zipper-like domain. J Biol Chem 280, 30185–30191.

47. Niwa, S., Tao, L., Lu, S.Y., Liew, G.M., Feng, W., Nachury, M.V., and Shen, K. (2017). BORC Regulates the Axonal Transport of Synaptic Vesicle Precursors by Activating ARL-8. Curr Biol 27, 2569–2578 e2564.

48. Ozkan, N., Koppers, M., van Soest, I., van Harten, A., Jurriens, D., Liv, N., Klumperman, J., Kapitein, L.C., Hoogenraad, C.C., and Farias, G.G. (2021). ER - lysosome contacts at a pre-axonal region regulate axonal lysosome availability. Nat Commun 12, 4493.

49. Pal, P., Taylor, M., Lam, P.Y., Tonelli, F., Hecht, C.A., Lis, P., Nirujogi, R.S., Phung, T.K., Yeshaw, W.M., Jaimon, E., et al. (2023). Parkinson’s VPS35[D620N] mutation induces LRRK2-mediated lysosomal association of RILPL1 and TMEM55B. Sci Adv 9, eadj1205.

50. Pu, J., Guardia, C.M., Keren-Kaplan, T., and Bonifacino, J.S. (2016). Mechanisms and functions of lysosome positioning. J Cell Sci 129, 4329–4339.

51. Pu, J., Keren-Kaplan, T., and Bonifacino, J.S. (2017). A Ragulator-BORC interaction controls lysosome positioning in response to amino acid availability. J Cell Biol 216, 4183–4197.

52. Pu, J., Schindler, C., Jia, R., Jarnik, M., Backlund, P., and Bonifacino, J.S. (2015). BORC, a multisubunit complex that regulates lysosome positioning. Dev Cell 33, 176–188.

53. Roney, J.C., Cheng, X.T., and Sheng, Z.H. (2022). Neuronal endolysosomal transport and lysosomal functionality in maintaining axonostasis. J Cell Biol 221.

54. Rudnik, S., Heybrock, S., Saftig, P., and Damme, M. (2022). S-palmitoylation determines TMEM55B-dependent positioning of lysosomes. J Cell Sci 135.

55. Saftig, P., and Puertollano, R. (2021). How Lysosomes Sense, Integrate, and Cope with Stress. Trends Biochem Sci 46, 97–112.

56. Sato, T., Ishikawa, M., Mochizuki, M., Ohta, M., Ohkura, M., Nakai, J., Takamatsu, N., and Yoshioka, K. (2015). JSAP1/JIP3 and JLP regulate kinesin-1-dependent axonal transport to prevent neuronal degeneration. Cell Death Differ 22, 1260–1274.

57. Scott, I., Webster, B.R., Chan, C.K., Okonkwo, J.U., Han, K., and Sack, M.N. (2014). GCN5-like protein 1 (GCN5L1) controls mitochondrial content through coordinated regulation of mitochondrial biogenesis and mitophagy. J Biol Chem 289, 2864–2872.

58. Snouwaert, J.N., Church, R.J., Jania, L., Nguyen, M., Wheeler, M.L., Saintsing, A., Mieczkowski, P., Manuel de Villena, F.P., Armao, D., Moy, S.S., et al. (2018). A Mutation in the Borcs7 Subunit of the Lysosome Regulatory BORC Complex Results in Motor Deficits and Dystrophic Axonopathy in Mice. Cell Rep 24, 1254–1265.

59. Song, A.H., Wang, D., Chen, G., Li, Y., Luo, J., Duan, S., and Poo, M.M. (2009). A selective filter for cytoplasmic transport at the axon initial segment. Cell 136, 1148–1160.

60. Stiernagle, T. (2006). Maintenance of C. elegans. WormBook, 1–11.

61. Takemasu, S., Nigorikawa, K., Yamada, M., Tsurumi, G., Kofuji, S., Takasuga, S., and Hazeki, K. (2019). Phosphorylation of TMEM55B by Erk/MAPK regulates lysosomal positioning. J Biochem 166, 175–185.

62. Trapnell, C., Roberts, A., Goff, L., Pertea, G., Kim, D., Kelley, D.R., Pimentel, H., Salzberg, S.L., Rinn, J.L., and Pachter, L. (2012). Differential gene and transcript expression analysis of RNA-seq experiments with TopHat and Cufflinks. Nat Protoc 7, 562–578.

63. Ungewickell, A., Hugge, C., Kisseleva, M., Chang, S.C., Zou, J., Feng, Y., Galyov, E.E., Wilson, M., and Majerus, P.W. (2005). The identification and characterization of two phosphatidylinositol-4,5-bisphosphate 4-phosphatases. Proc Natl Acad Sci U S A 102, 18854–18859.

64. Verhey, K.J., Meyer, D., Deehan, R., Blenis, J., Schnapp, B.J., Rapoport, T.A., and Margolis, B. (2001). Cargo of kinesin identified as JIP scaffolding proteins and associated signaling molecules. J Cell Biol 152, 959–970.

65. Vilela, F., Chenon, M., Campanacci, V., Alik, A., Velours, C., Andreani, J., Llinas, P., and Menetrey, J. (2026). Insights into how JIP3 and JIP4 adaptor proteins relieve the auto-inhibited form of the kinesin-1 motor. J Biol Chem, 113355.

66. Vilela, F., Velours, C., Chenon, M., Aumont-Nicaise, M., Campanacci, V., Thureau, A., Pylypenko, O., Andreani, J., Llinas, P., and Menetrey, J. (2019). Structural characterization of the RH1-LZI tandem of JIP3/4 highlights RH1 domains as a cytoskeletal motor-binding motif. Sci Rep 9, 16036.

67. Wallings, R.L., Humble, S.W., Ward, M.E., and Wade-Martins, R. (2019). Lysosomal Dysfunction at the Centre of Parkinson’s Disease and Frontotemporal Dementia/Amyotrophic Lateral Sclerosis. Trends Neurosci 42, 899–912.

68. Wang, C., Telpoukhovskaia, M.A., Bahr, B.A., Chen, X., and Gan, L. (2018). Endo-lysosomal dysfunction: a converging mechanism in neurodegenerative diseases. Curr Opin Neurobiol 48, 52–58.

69. Waschbusch, D., Pal, P., Nirujogi, R.S., Cavin, M., Singh, J., Alessi, D.R., and Khan, A.R. (2026). Structural basis for binding of RILPL1 to TMEM55B reveals a lysosomal platform for adaptor assembly through a conserved peptide motif. Structure 34, 296–310 e295.

70. Watt, D., Dixit, R., and Cavalli, V. (2015). JIP3 Activates Kinesin-1 Motility to Promote Axon Elongation. J Biol Chem 290, 15512–15525.

71. Wen, Q., Po, M.D., Hulme, E., Chen, S., Liu, X., Kwok, S.W., Gershow, M., Leifer, A.M., Butler, V., Fang-Yen, C., et al. (2012). Proprioceptive coupling within motor neurons drives C. elegans forward locomotion. Neuron 76, 750–761.

72. Willett, R., Martina, J.A., Zewe, J.P., Wills, R., Hammond, G.R.V., and Puertollano, R. (2017). TFEB regulates lysosomal positioning by modulating TMEM55B expression and JIP4 recruitment to lysosomes. Nat Commun 8, 1580.

73. Wilm, T., Demel, P., Koop, H.U., Schnabel, H., and Schnabel, R. (1999). Ballistic transformation of Caenorhabditis elegans. Gene 229, 31–35.

74. Winckler, B., Faundez, V., Maday, S., Cai, Q., Guimas Almeida, C., and Zhang, H. (2018). The Endolysosomal System and Proteostasis: From Development to Degeneration. J Neurosci 38, 9364–9374.

75. Yap, C.C., Digilio, L., McMahon, L.P., Garcia, A.D.R., and Winckler, B. (2018). Degradation of dendritic cargos requires Rab7-dependent transport to somatic lysosomes. J Cell Biol 217, 3141–3159.

76. Zahavi, E.E., Hummel, J.J.A., Han, Y., Bar, C., Stucchi, R., Altelaar, M., and Hoogenraad, C.C. (2021). Combined kinesin-1 and kinesin-3 activity drives axonal trafficking of TrkB receptors in Rab6 carriers. Dev Cell 56, 494–508 e497.

77. Zhang, A., He, X., Zhang, L., Yang, L., Woodman, P., and Li, W. (2014). Biogenesis of lysosome-related organelles complex-1 subunit 1 (BLOS1) interacts with sorting nexin 2 and the endosomal sorting complex required for transport-I (ESCRT-I) component TSG101 to mediate the sorting of epidermal growth factor receptor into endosomal compartments. J Biol Chem 289, 29180–29194.

78. Zhou, B., Zhu, Y.B., Lin, L., Cai, Q., and Sheng, Z.H. (2011). Snapin deficiency is associated with developmental defects of the central nervous system. Biosci Rep 31, 151–158.

