## Supplemental Figures for "TMEM55B promotes anterograde transport and distal axonal distribution of LAMP1-positive organelles"

### Supplemental Figure 1

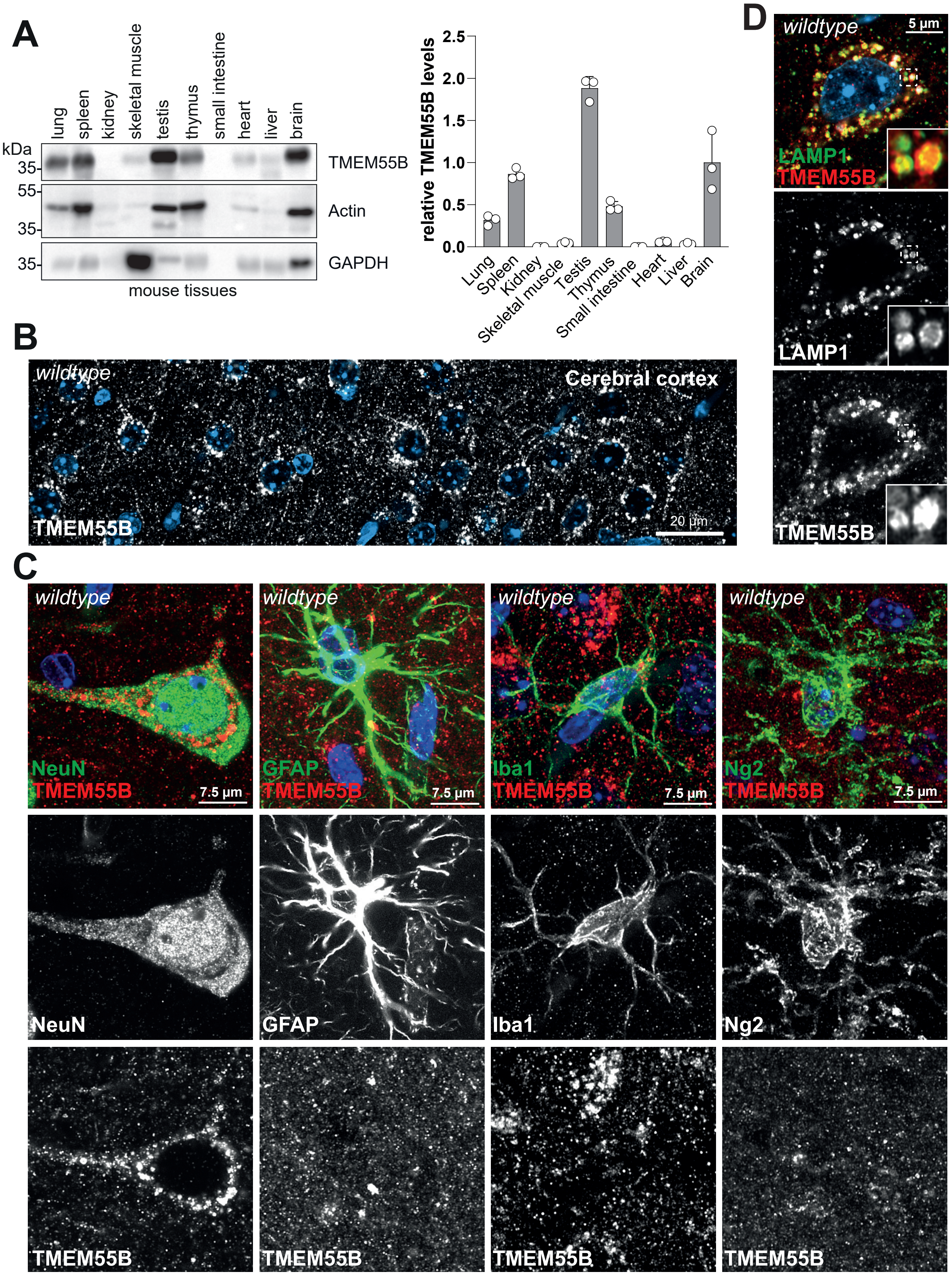

### Supplemental Figure 2

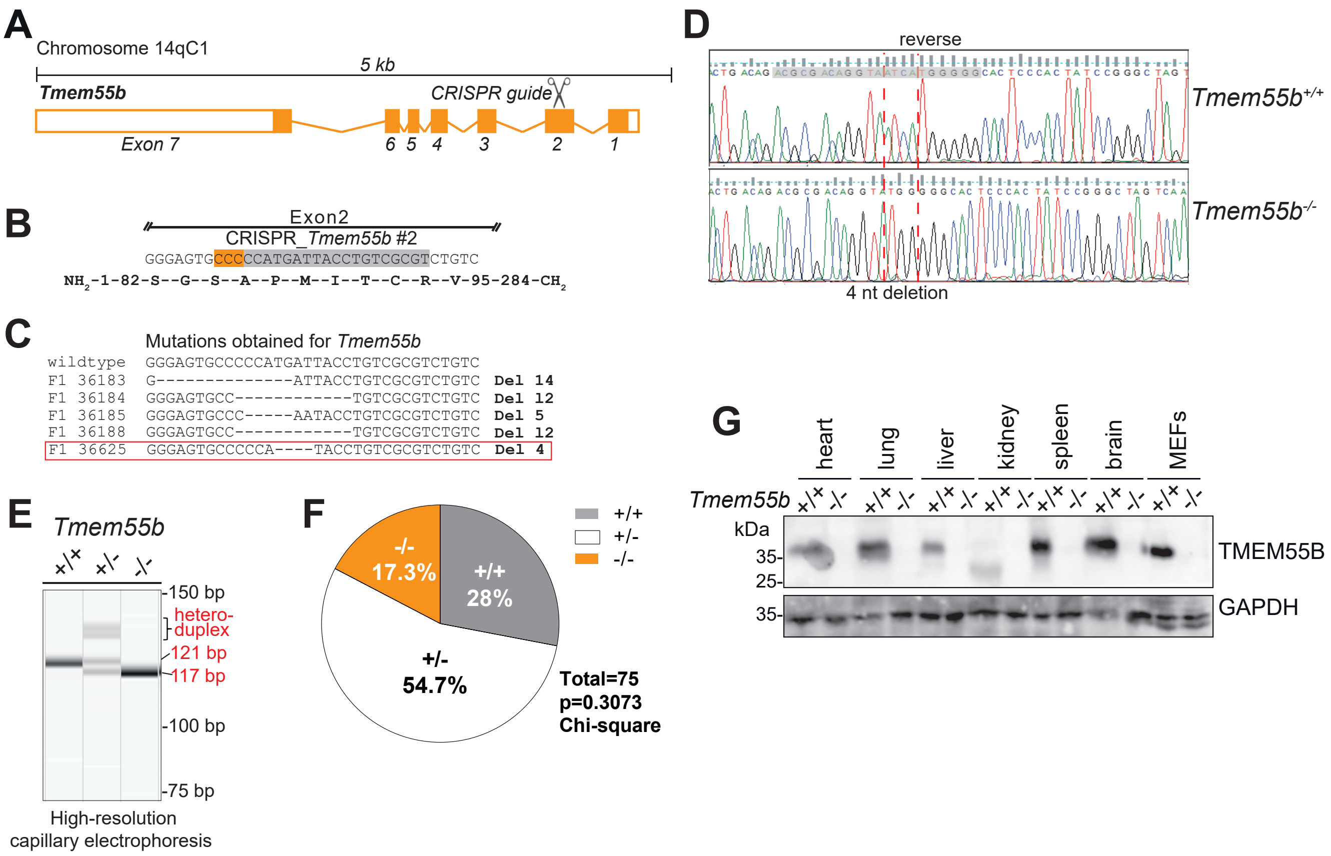

#### Supplemental Figure 3

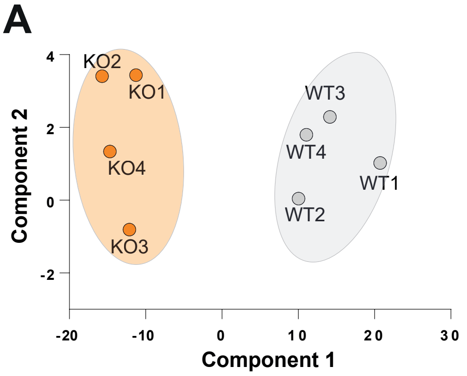


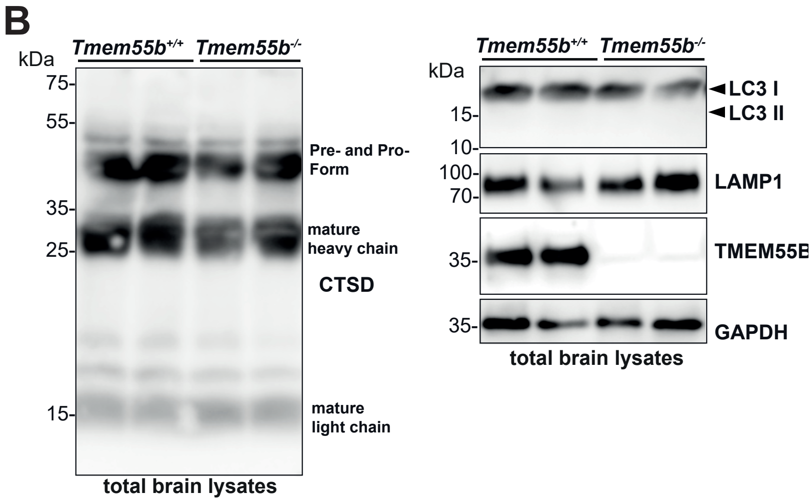

### Supplemental Figure 4

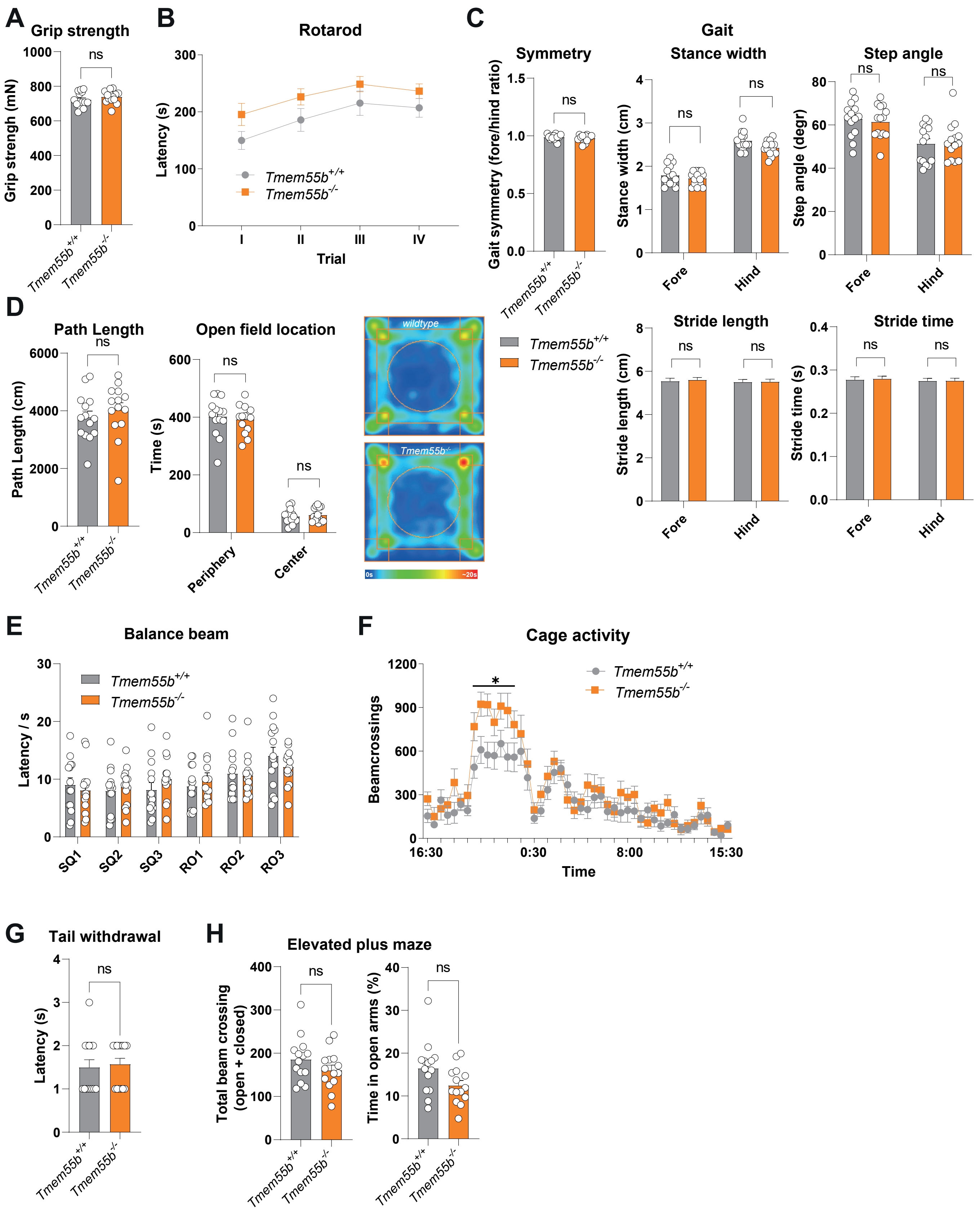

### Supplemental Figure 5

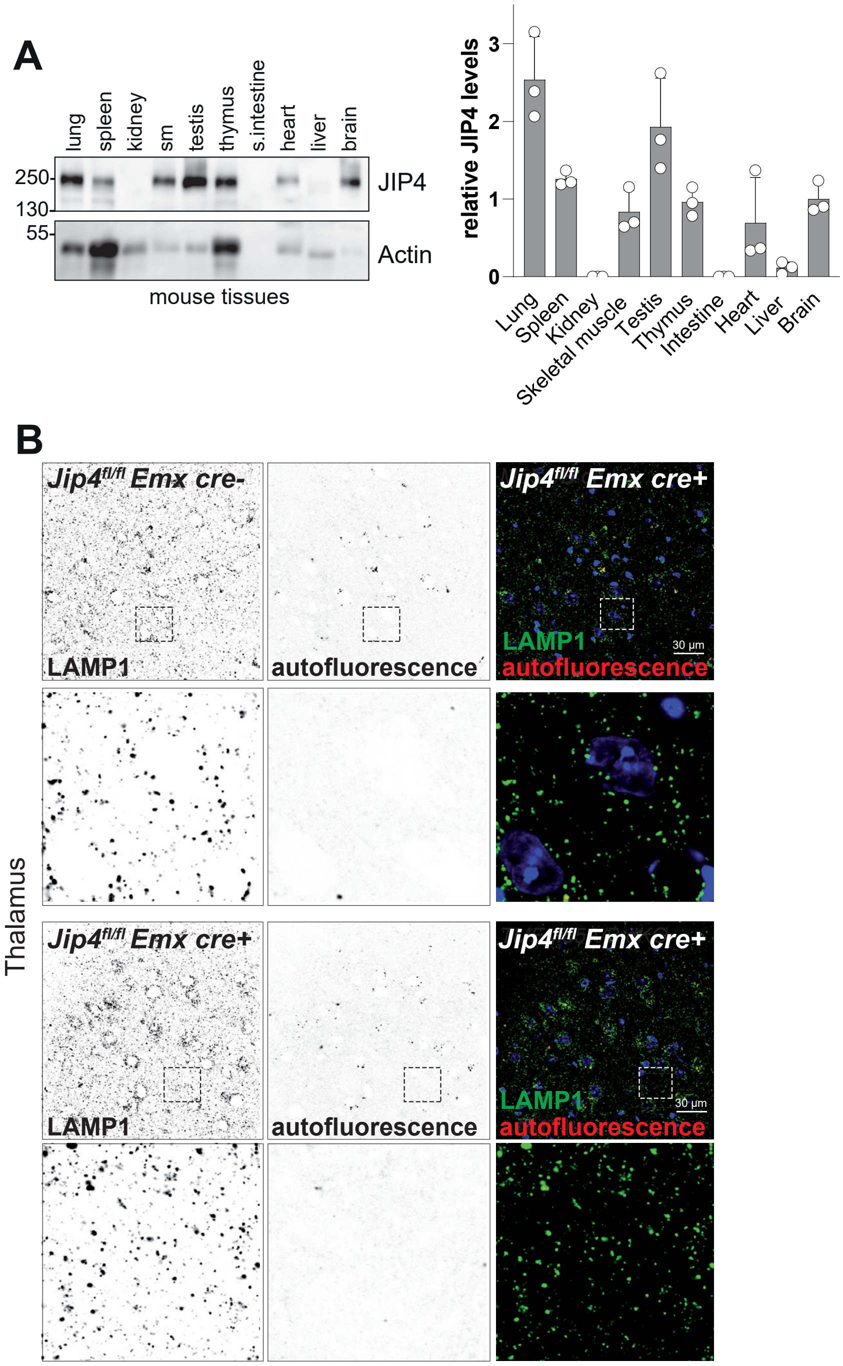

### Supplemental Figure 6

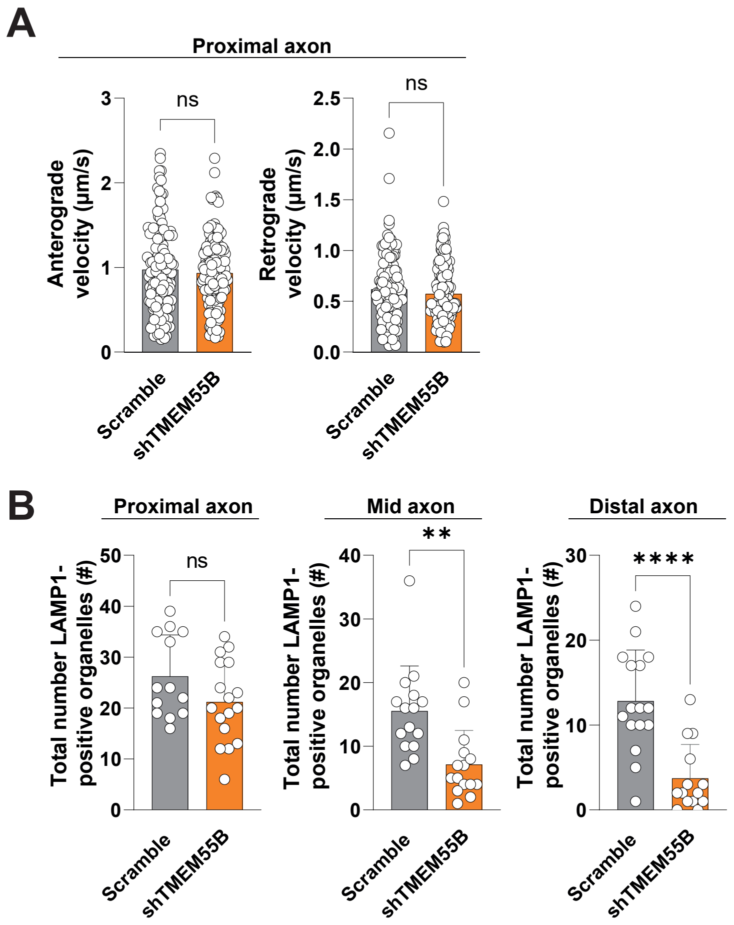

### Supplemental Figure 7

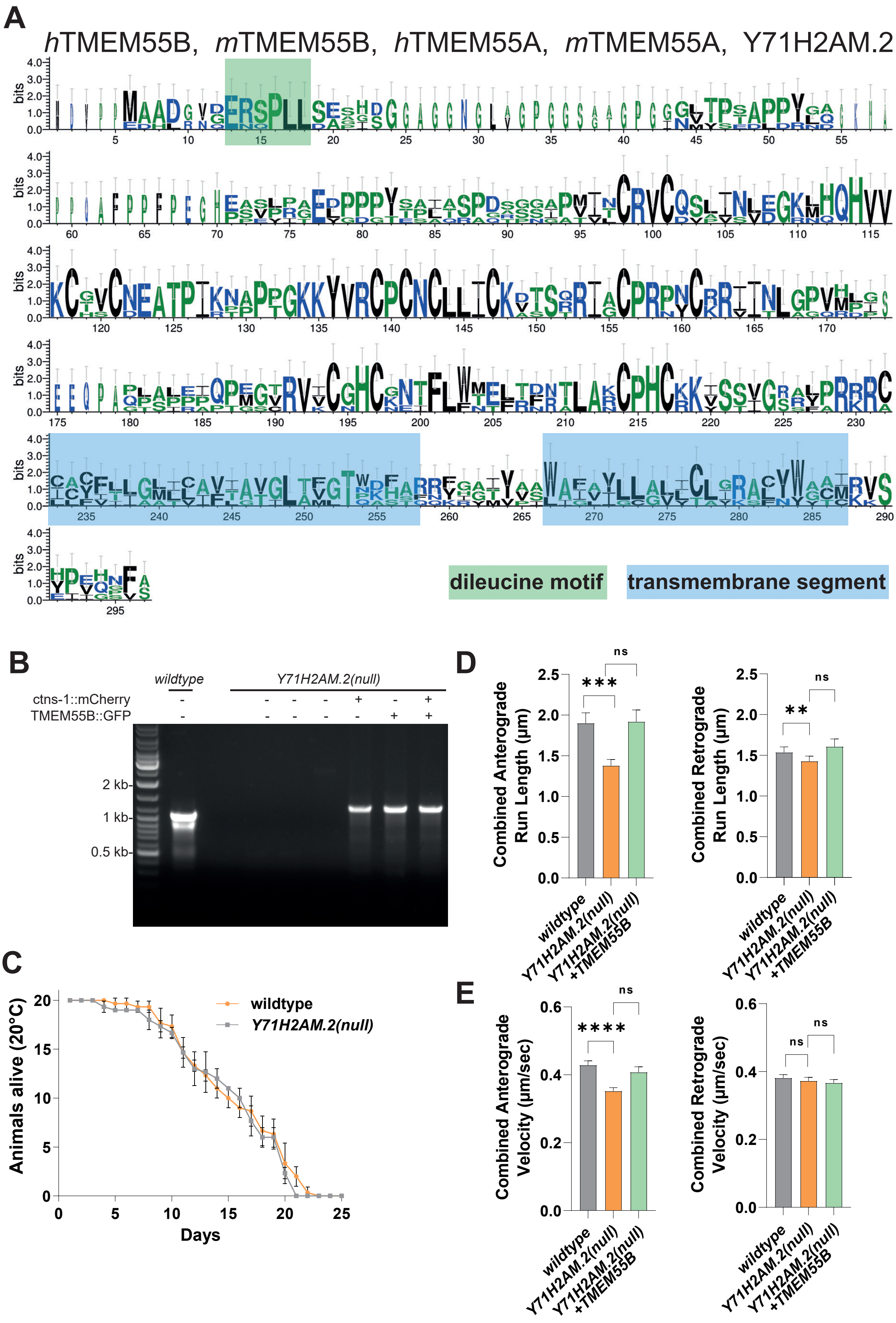

### Supplemental Figure 8

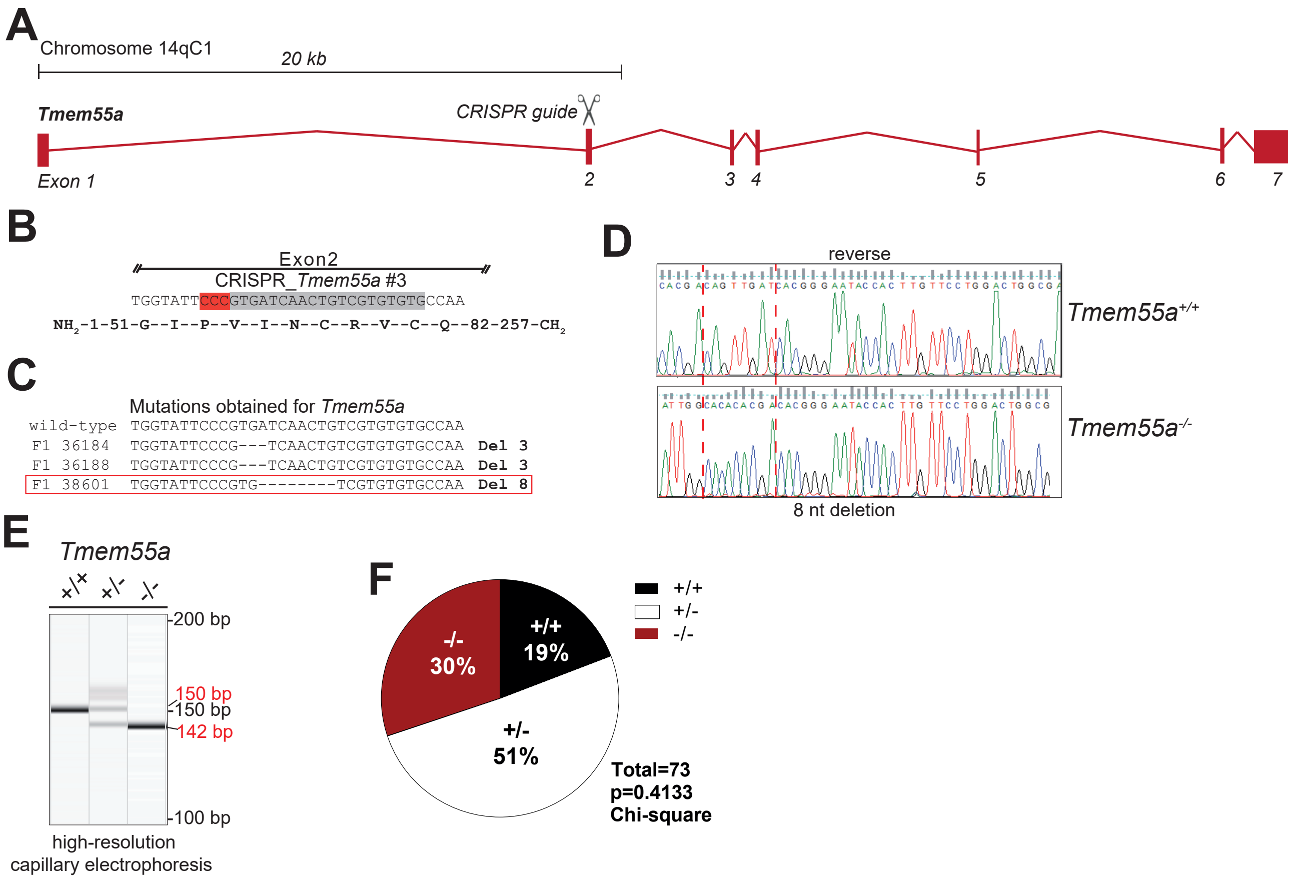
