## Supplementary figures and images for "TMEM55B promotes anterograde transport and distal axonal distribution of LAMP1-positive organelles"

### Scramble C2-20241029 DIV8Hippo LAMP1GFP Neuron3_w1561_t1 Soma Zoom.tif

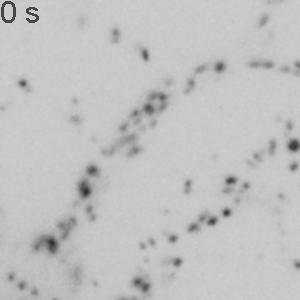

### shTMEM55B1_2_C2-20241029 DIV8Hippo LAMP1GFP Neuron2_w1561_t1 Soma zoom.tif

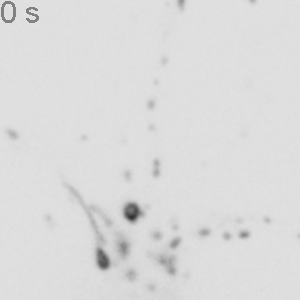
